# The broad ecological range of *Salmonella enterica* serotype Typhimurium is associated with genomic diversification

**DOI:** 10.64898/2026.04.16.719076

**Authors:** Leela Ohri, Sandeep Chinnareddy, Ying-Xian Goh, Hailong Zhang, Xiangyu Deng, Amy Pruden, Rachel Cheng, Song Li, Jingqiu Liao

## Abstract

*Salmonella* Typhimurium is a versatile foodborne pathogen with a broad ecological range, making it an ideal model to better understand pathogen adaptations that allow them to infect multiple hosts and persist across environments. Here, we analyzed 595 genomes of *S.* Typhimurium representing three food animal sources (bovine, swine, and poultry) and one non-food animal source (wild birds). We found that *S.* Typhimurium from food animal sources generally had a more open pangenome and harbored more antimicrobial resistance genes (ARGs). Notably, swine isolates exhibited the most open pangenome and the highest prevalence of ARGs, patterns that were associated with a greater presence of mobile genetic elements, particularly plasmids. Despite similar core genome sizes, *S.* Typhimurium from different sources displayed distinct patterns of positive selection in the core genome that varied in frequency and targeted functional categories. In contrast, although accessory genome sizes varied substantially across sources, the frequency of positive selection remained similar. Using machine learning, we identified source-predictive genetic variants, many of which are associated with stress-response functions. These findings suggest that gain and loss of accessory genes and positive selection acting on core genes support differential adaptation in *S.* Typhimurium, potentially contributing to its broad ecological range.

## 1 Introduction

Some pathogens can infect a wide range of hosts; including humans, animals, and plants, and persisting across diverse niches such as soil, water, and built environments^1–4^. Such ecological versatility bears particular importance for foodborne pathogens, as their ability to withstand diverse stressors and transmit across ecosystem boundaries contributes to their persistence along the farm-to-fork continuum, posing major challenges to food safety and public health^5–9^. As such, dissecting the genomic traits that underpin the broad host and environmental adaptability of foodborne pathogens is vital not only for advancing the mechanistic understanding of bacterial adaptive evolution, but also for improved predictive surveillance, risk assessment, and targeted interventions to safeguard the food supply.

Bacterial genomes often exhibit a high degree of flexibility, characterized by the corresponding pangenome, which refers to the total repertoire of genes pertaining to a given species or clade of interest^10^ ^11^. Some species have a more closed pangenome with many genes overlapping among individual genomes (i.e., core genes). On the other hand, other species have a more open pangenome, harboring a high proportion of genes not shared between individual genomes (i.e., accessory genes)^11,12^. These accessory genes often allow for specialization of traits such as pathogenicity and antimicrobial resistance (AMR) amongst different strains and organisms^13^. The openness of a pangenome, operationally defined as the rate of expansion as more genomes are sequenced, has been linked to adaptive evolution, reflecting the capability of a population to adjust to changing environments^14^. Gene gain and loss drive genome evolution through multiple mechanisms, including the formation of ‘pseudogenes’ via point mutations as well as horizontal gene transfer (HGT), which involves the impartment of genetic information between organisms that are not parent-offspring^15–17^. HGT can occur through transformation, transduction, and conjugation^16^. Enabling almost immediate sharing of genetic information, HGT enables bacteria to rapidly respond to environmental stressors^18^. A prominent example is the dissemination of antimicrobial resistance genes (ARGs) across the food production chain, which poses an urgent threat to disease control, including increased morbidity and mortality and higher healthcare costs^19–21^. While ARGs can arise internally through spontaneous mutation, their widespread dissemination is largely driven by external acquisition via HGT, a process that can be accelerated by the overuse and misuse of antibiotics in the agriculture and human healthcare sectors^22–25^.

Positive selection, in which genetic mutations or gene variants that confer a beneficial trait are favored and increase in frequency within a bacterial population over time,^26^ is also a fundamental driver of bacterial adaptive evolution. Both core and accessory genomes are subject to positive selective pressure, though it may manifest differently. Bacterial core genomes primarily undergo a form of purifying (or negative) selection, eliminating detrimental genes or mutations to conserve energy and resources, creating highly conserved genomes^27,28^. Accessory genomes, which are subjected to higher rates of change, often experience stronger positive selection. With the introduction of novel genes into individual genomes, those that are beneficial to the organism’s survival and reproductive capabilities, become widespread while those that are neutral or detract are discarded^28^. Of the genes that do undergo positive selection, the functions are varied but often include ARGs and those coding for cell surface proteins – which can be immune-response and phage predation targets, and carbon acquisition pathways^29–32^. Foodborne pathogens often encounter high-stress conditions (e.g., nutrient limitation, temperature extremes, oxidative stress, chemical stress, and osmotic stress) throughout the food production continuum. Mutations can sometimes confer tolerance to such stressors and thus be subject to positive selective pressure and potential to be permanently fixed in the bacterial population^9^.

As one of the primary agents of foodborne illness, *Salmonella enterica* causes an estimated 1.35 million infections annually in the United States (US), resulting in substantial hospitalizations and fatalities. *S. enterica* is genetically diverse, possessing over 2,600 serovars and at least six subspecies^33–35^. One of the non-typhoidal serotypes (NTS) of *S. enterica*, *S.* Typhimurium, is known to interact with a broad range of hosts, such as swine, cows, poultry animals, and wild birds^36–38^. Of note, AMR is emerging in *S.* Typhimurium^39^, with many multidrug-resistant (MDR) strains showing a high degree of resistance towards commonly prescribed antibiotics^40^. For example, one study of 277 *S.* Typhimurium isolates collected in China over the course of 10 years (2007-2019) identified MDR strains in 77.3% of isolates and resistance to ciprofloxacin in 11.6%^41^. Another study based on Iranian poultry markets yielded isolates of which 72% contained resistance to tetracyclines and over half of all detected isolates demonstrated some form of resistance^42^. AMR in *S.* Typhimurium is maintained and disseminated across the food production chain. Proximal location creates an element of concern, with nearby farms transmitting resistance across space and species^43^, passing from poultry to cattle and vice versa. Importantly, many of these ARGs are on mobile genetic elements (MGEs)^44^, mainly plasmids, transposable elements, or bacteriophages^45^, facilitating the emergence and dissemination of AMR in this pathogen via HGT. Additionally, evidence of positive selection has been identified in *S.* Typhimurium, such as on genes encoding cell surface proteins and ARGs^32,46^. Given its high ecological versatility, genomic flexibility, and public health relevance, *S.* Typhimurium presents itself as an ideal candidate for addressing gaps in our understanding of the genomic bases underpinning the broad ecological niches of foodborne pathogens.

Here, we performed comparative genomics of 595 high-quality *S.* Typhimurium genomes representing four broad sources, including three food animal sources - bovine, poultry, and swine, and one non-food animal source - wild birds. To identify source-associated variation in the core and accessory genomes and evolutionary processes, we compared pangenome structure, positive selection frequency, and MGE and ARG profiles across isolation sources. We also developed machine learning models to predict isolation sources based on patterns of gene presence/absence and core single nucleotide polymorphisms (SNPs). Overall, this study provides valuable insights into possible drivers of adaptive evolution of *S.* Typhimurium and potential genomic mechanisms underlying its broad ecological range. In addition, our machine learning-based predictive models can support efficient source investigation of *S.* Typhimurium by identifying associated genomic features, enhancing food safety decision-making.

## 2 Experimental Procedures

### 2.1 *S.* Typhimurium genomic data

*S.* Typhimurium genome assemblies collected in the US and uploaded in 2021 were retrieved from the National Center for Biotechnology Information (NCBI) Pathogen Detection database (collection dates ranging 2002-2021), yielding a total of 745 assemblies. The year of 2021 was chosen because it was the most recent year at the time this analysis was initiated (**Table S1**). Isolates were selected by upload date rather than collection date because collection-year metadata was unavailable for many genomes. This approach allowed us to retain a larger dataset. Based on the isolation sources, these genomes were classified into five major sources, including three food animal sources (bovine, poultry, and swine) and two non-food animal source (wild bird and the environment); isolation source categorization was done using the ‘isolation_source’ field information provided in the NCBI Pathogen Detection database. For food animal sources, the common or Latin name (for example: *Bos taurus* for cow) of the animal or animal part (e.g., kidney, raw intact beef) was used to assign the category. Wild birds included any bird except turkey or chicken (these were collectively classified as ‘poultry’). As the environmental category is broad and is composed of distinct subcategories (e.g., water, tree nut, water filtration membrane, litter, and soil), it is warranted to comprehensively assess the genomic diversification of *S.* Typhimurium across environmental subcategories, rather than grouping them into a single broad category, to better understand patterns of environmental adaptation and evolution. Therefore, only bovine, swine, poultry, and wild bird remained as the sources examined in the study, reducing the genome sample set size to 654. To assess the quality of the assemblies, QUAST v.4.0^47^, CheckM2^48^, Kraken2^49^, and SISTR^50^ were employed with the default settings for each program. All the assemblies passed the following quality controls thresholds: number of contigs < 300 (mean ± sd of 105.19 ± 57.00) and an N50 > 20,000 (mean ± sd of 272,948.95 ± 774,636.37), completeness > 90% (99.99 ± 0.0016), contamination < 5% (3.28 ± 1.28), taxonomic identification as *Salmonella*, and a positive serotype identification as *Salmonella enterica* serotype Typhimurium (**Table S1**). A total of 59 assemblies did not meet the quality criteria, and were thus excluded from downstream analyses. The final data set included 595 isolates, including 61 bovine, 285 poultry, 56 swine, and 193 wild bird isolates. The genome size ranged between 4.72 Mbp to 5.38 Mbp (mean ± sd of 4.96 ± 0.088 Mbp), which is an expected genome size for *S. enterica*^51^.

### 2.2 Gene prediction, orthologous gene identification, functional annotation, core SNP identification, and phylogenetic tree construction

Genes in each genome were predicted using Prodigal v.2.6.3^52^; pseudogenes were not annotated and were treated as gene absence in order to characterize the repertoire of potentially functional genes rather than all homologous sequence remnants. This choice may lead to a more conservative estimate of accessory gene presence, particularly for genes that have undergone recent loss of function. Orthologous genes across genomes were identified using MMseqs2^53^ with a coverage of 0.8 and a minimal identity of 0.5. These cutoffs were chosen to allow detection of homologous genes that may have accumulated substantial sequence divergence while still requiring most of the sequence length to align, reducing artificial splitting of divergent homologs into separate gene families. However, a relatively permissive identity threshold may merge more distantly related homologs and therefore affect estimates of core and accessory genome size. Amino acid sequences of the representative gene for each orthologous gene were functionally annotated using eggNOG v.5.0^54^. Amino acid sequences of each orthologous gene were aligned using MUSCLE v.3.8.31^55^ and were converted to codon alignments using PAL2NAL v.14^56^ with gaps removed. Homologous recombination was assessed using 3SEQ v.1.8.0^57^ and the MAXCHI algorithm^58^ implemented in PHIPACK v.1.1. Only recombination events detected by both methods were considered evidence of homologous recombination. Non-paralogous, non-recombinant core genes (i.e., orthologous genes present in all 595 *S.* Typhimurium genomes and having no paralogs as well as no evidence of homologous recombination) were concatenated. Core SNPs were identified based on these core genes using a custom Python script^14^. A maximum-likelihood phylogenetic tree of the 595 *S.* Typhimurium genomes was reconstructed from the core SNP alignment using IQ-TREE v.3.0.1^59^ with 1,000 bootstraps. The best-fitting evolutionary model (TVM+F+ASC+R3) was selected using ModelFinder^60^, implemented in IQ-TREE v.3.0.1, based on Bayesian information criterion (BIC). The tree, rooted by mid-point, was visualized using iTOL v.7.6^61^. To assess if clusters of isolates exist, lineage models were fitted to the genomes using PopPUNK v2.7.0, a software widely used for population analysis and clustering, at Ranks 1, 2, 3, 5, and 10^62^. Rank refers to the resolution of the model, with Rank = 1 being the most specific.

### 2.3 Pangenome and core genome characterization

The pangenome of a *S.* Typhimurium population for each source was defined as the entire sets of orthologous genes for individuals isolated from the same source. The core genome was defined as the orthologous genes found in all individuals isolated from the same source, while the accessory genome was defined as the orthologous genes not shared by all individuals. Rarefaction curves of pangenome and core genome of each population from a given source were estimated by subsampling an increasing number of genomes using an R script provided by Méric et al (2014)^63^. The pangenome and core genome curves were fit to the power law function *cN^γ^*, where *c* is the pangenome or core genome size when *N* is 1, *N* is the number of genomes, and *γ* is a scaling exponent ranging between 0 and 1. The pangenome and core genome size of each population given *N* of 50 was predicted using the fitted power law function.

Principal Coordinates Analysis (PCoA) of the presence/absence matrix of accessory genes calculated using the Jaccard distance were performed to compare accessory genome composition for populations with different isolation sources, followed by permutational multivariate analysis of variance (PERMANOVA) and permutational multivariate analysis of dispersion (PERMDISP) tests.

### 2.4 Plasmid, prophage, and AMR profiling

Plasmids were predicted using Platon in the accuracy mode and filtered for a Replicon Distribution Score (RDS) > 0.7, indicating a confident classification of a contig as plasmid-derived rather than chromosomal^64^. Prophages were predicted using PHASTEST with default settings^65^. ARGs were identified using BLASTN searches^66^, with an E-value of 0.01 and without restrictions on percent identity, against the Comprehensive Antibiotic Resistance Database (CARD)^67^. The best match of each ARG was chosen based on the highest bit-score and assessed for the presence of premature stop codons (TGA, TAG, TAA; up to but not including the last stop codon^68^) and sequence coverage (%). ARGs were categorized as “putatively functional” if their sequence coverage exceeded 80% and no premature stop codon was detected; truncated if its sequence coverage ranged between 30% - 80% or a premature stop codon was detected; and absent if sequence coverage was less than 30% or no hits were observed in the BLASTN searches according to the criteria detailed in Liao et al (2023)^69^. Based on this categorization, the classification of ARGs was further simplified as “present ARGs” (including either truncated or putatively functional) and “functional ARGs”. Here, the term “functional ARGs” denotes the presence of sufficient genetic sequence to encode a functional protein upon expression; it does not imply the active transcriptomic state of the bacterium. CARD-RGI v.6.0.8^70^ was used to validate our ARG prediction. A comparison was subsequently conducted to determine if the functional or present ARGs were found on plasmids and prophages based on their genomic coordinates. Notably, although Platon improves plasmid-chromosome discrimination by integrating RDS scores, it operates on fragmented contigs generated from short-read assemblies^64^. Also, PHASTEST could over- or under-estimate prophage presence, as draft assemblies can cause either artificial duplication or fragmentation^71^. As such, ARGs identified on contigs predicted to be derived from plasmids or prophages should be considered putatively MGE-associated until their genomic contexts are confirmed using long-read or hybrid assemblies or an independent validation pipeline.

To assess the differences in the sequence proportion of plasmids and prophages, and count of total and unique present and functional ARGs among isolation sources, Kruskal– Wallis tests were conducted followed by Mann-Whitney *U* tests for pairwise comparisons. Fisher’s exact tests were performed to assess the differences in the frequency of individual ARGs, their drug classes, and families among isolation sources followed by Benjamini-Hochberg (BH) false discovery rate (FDR) correction to account for multiple testing. Kruskal–Wallis tests were performed to assess the differences in the count of ARGs belonging to each resistance mechanism compared among isolation sources followed by BH FDR correction.

To account for possible biases due to differing sample sizes across isolate sources, a parallel ARG and MGE analysis was conducted utilizing a subsampled data set with N = 56 isolates randomly chosen from each isolate source. To determine the possible effect of a temporal bias on the data, a subsampled set of isolates both collected and uploaded in 2021 was analyzed to assess trend similarity to the overall results of the study. The year 2021 was chosen due to offering the highest number of samples collected and uploaded, maximizing statistical potential for subsequent analyses.

### 2.5 Positive selection and functional enrichment analysis

The BUSTED-[S] model implemented in HyPhy^72,73^ was employed to test gene-wide positive selection in core and accessory genes for each population. Only non-paralogous, non-recombinant genes with more than three nonidentical sequences were included in the BUSTED analysis. A likelihood ratio test was conducted by comparing the unconstrained model allowing for positive selection against the constrained model disallowing for positive selection. Statistical significance was determined by approximating the likelihood ratio test statistic to a *χ*^2^ distribution. The differences in the frequency of all, core, and accessory genes under positive selection were compared among isolation sources using Fisher’s exact tests.

To identify COG (Clusters of Orthologous Groups) functional categories that were significantly overrepresented among genes showing evidence of positive selection, a binomial framework was used to compare the frequency of each COG among genes under positive selection against its genome-wide frequency as the null expectation using the formula below:

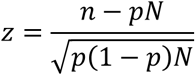

where *n* is the observed count of genes showing evidence of positive selection that belong to a COG, *N* is the total count of genes, *p* is the frequency of genes belonging to a COG among all genes, and *z* is the standardized binomial Z-score, which represents the multiplier of standard deviation in the binomial distribution. COGs with a standardized binomial Z-score greater than or less than -2 were considered significantly enriched or depleted, respectively, corresponding approximately to a two-sided *P* < 0.05 under the normal approximation. The functional enrichment analysis was performed for both core and accessory genes for each *S.* Typhimurium population from a given source.

### 2.6 Machine learning models

To identify genetic variance that can be used to predict the isolation sources of *S.* Typhimurium, we developed a machine learning-based framework using isolation sources as the outcome and gene presence/absence or core SNPs as features. Samples collected from 2021 and those lacking a noted collection year (as most wild bird isolates did not have a known collection year) were first cleaned and split into the training set (80%) and the testing set (i.e., the holdout set; 20%) in a stratified fashion. This preserved the relative class proportion across folds and reduced potential bias associated with unequal sample sizes among source categories. The training set was further split into 5 stratified folds for cross-validation, in which a collection of predefined models (i.e., classifiers based on decision trees, random forest, multilayer perception, support vector machines [SVM], and gradient boosting) were trained and tested with a random set of hyperparameters. The average area under the receiver operating characteristic curve (auROC) score was used to evaluate the performance of the models across the 5 rounds of cross-validation. To account for stochasticity introduced by the random splitting of samples and division of training data into 5 folds, we repeated these steps 10 times. We selected the best model and its hyperparameter set with the highest interquartile mean of the auROC scores out of the 10 repetitions among the predefined models. The interquartile means of the auROC and area under the precision-recall curve (auPR) scores of the best model that was exclusively trained on the training set were reported based on a single evaluation of the holdout data from each of the 10 repetitions. The importance of the features (i.e., genes or core SNPs) was quantified using Shapley Additive exPlanations (SHAP)^74^. To ensure the models’ ability to predict accurately, despite temporal variability, the ML models with best performance were further validated on genomes with known collection years ranging from 2002 to 2020. This form of validation forces the model to predict across substantial temporal and genomic divergence.

## 3 Results

### 3.1 Pangenome composition in *S.* Typhimurium differs across isolation sources, with swine-derived isolates displaying the most open pangenome

To examine how pangenome composition varies by isolation source in *S.* Typhimurium, we analyzed the core and accessory genes of bovine, poultry, swine, and wild bird isolates. First, a phylogenetic tree was constructed based on core SNPs (**Fig. 1a**). We observed some phylogenetic intermixing among swine and bovine isolates, while wild bird and poultry isolates formed major clades predominantly composed of isolates from the same source. To further determine if clusters of isolates exist, we fitted lineage models to the genomes. Host-associated genomic differentiation was detectable primarily at fine clustering resolution (Rank 1 and Rank 2), whereas broader lineage definitions merge these host-associated sublineages into a common population (**Fig. S1**). The persistence of wild-bird-specific clusters at the finer ranks suggests that wild-bird-associated populations may contain genetically distinct sublineages. The observed clustering of isolates suggest an association between core genomes and isolation source in *S.* Typhimurium, particularly from wild birds.

**Figure 1:**
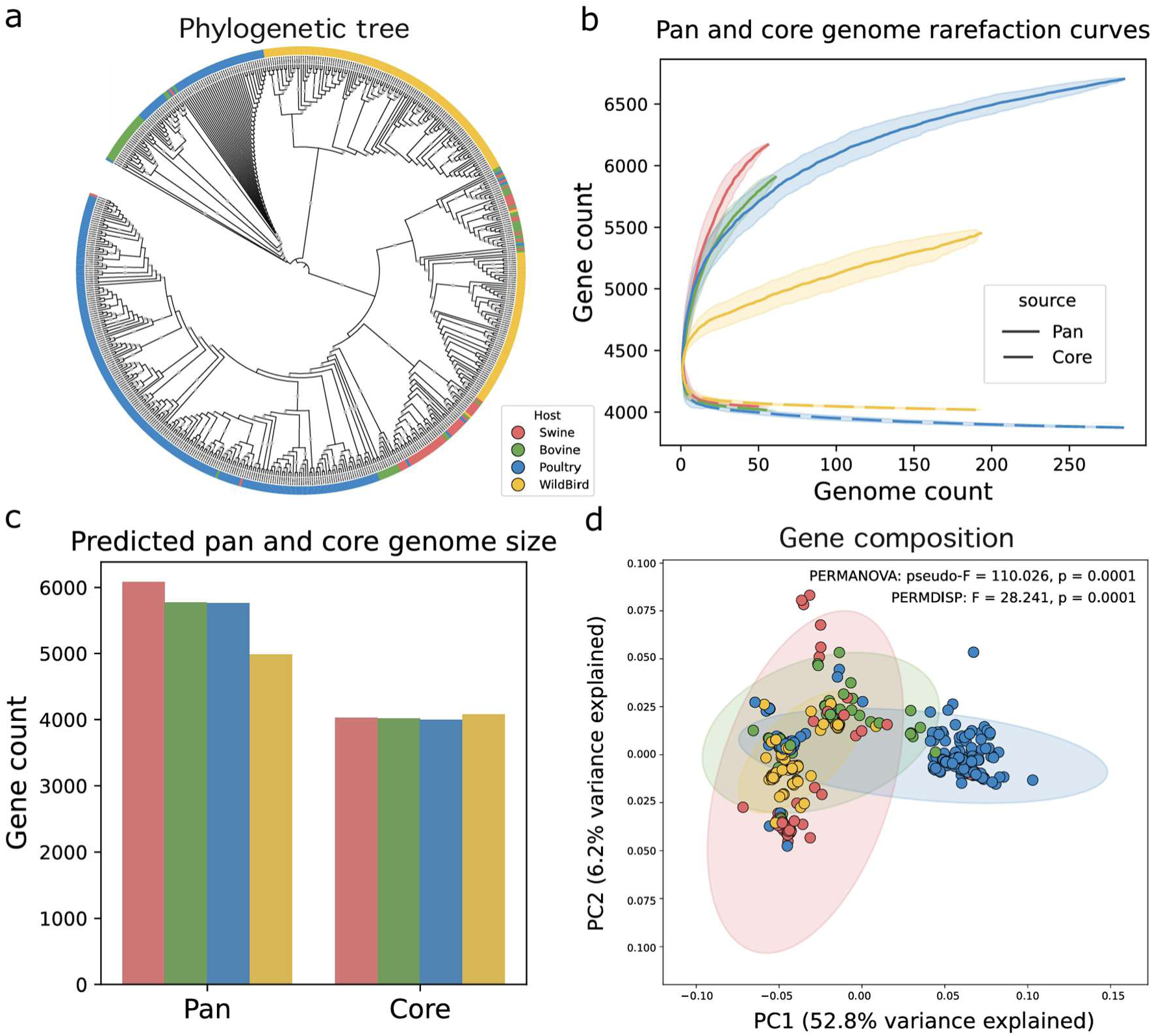
Pan genome structure of *S*. Typhimurium by isolation source. (a) A maximum likelihood phylogenetic tree of 595 *S*. Typhimurium genomes. The tree was constructed based on core SNPs with 1,000 bootstraps. Bootstrap values > 80% are indicated by grey circles. Tree tips are color coded by isolation sources (swine - red, bovine - green, poultry - blue, and wild bird - yellow). (b) Pan and core genome rarefaction curves for *S.* Typhimurium by isolation source. The colored lines and shaded areas depict the means and standard deviations, respectively, based on 100 randomized repetitions varying the order of genomes. The slope of the pangenome rarefaction curve reflects the degree of pangenome openness. (c) Bar plots showing the pan and core genome size of *S.* Typhimurium from different isolation sources given 50 genomes. The predicted pan and core genome sizes are derived from power-law functions fitted to the observed pangenome and core-genome curves for each population shown in (b). (d) PCoA plot showing gene composition differences of *S.* Typhimurium by isolation source. Each dot corresponds to one genome, with proximity of dots indicating similarity in gene composition. PERMANOVA: permutational multivariate analysis of variance; PERMDISP: permutational multivariate analysis of dispersion.

Next, we compared the pangenome and core genome sizes of *S.* Typhimurium populations across different isolation sources. Rarefaction curves showed that core genome size stabilized at approximately 3,800 genes, showing minimal sensitivity to the number of genomes analyzed, with populations from all sources exhibiting comparable levels of similarity (**Fig. 1b**). In contrast, the pangenomes showed wider variability across sources (**Fig. 1b**). The swine isolates exhibited the steepest pangenome curve, indicating the most open pangenome, followed by poultry and bovine isolates, which showed similar levels of pangenome openness. Conversely, indicated by a flatter curve, the pangenomes of wild bird isolates tend to be more closed. To account for potential bias caused by unbalanced sample sizes for different isolation sources, we fitted each pangenome and core genome curve to a power law function (see **Methods**) and predicted the pangenome and core genome sizes for each isolation source at an equal sample size of 50 genomes, which is the largest multiple of ten below the minimum number of genomes in any source category (**Fig. 1c**). As expected, core genome sizes were similar among populations from different sources, with 4,026, 4,015, 3,991, and 4,073 genes for swine, bovine, poultry, and wild bird isolates, respectively. In contrast, predicted pangenome size varied substantially across sources: swine isolates had the largest pangenome (6,082 genes), followed by bovine (5,771), poultry (5,759), and wild bird (4,980).

The differences in pangenome structure among isolation sources were further supported by the PCoA analysis, in which genomes formed clusters by isolation source based on gene presence/absence profiles (**Fig. 1d**), Statistical tests showed that gene presence/absence profiles differed significantly among isolation source groups (PERMANOVA *P* < 0.05). However, dispersion also differed significantly among groups (PERMDISP *P* < 0.05), indicating that the PERMANOVA result may reflect both differences in pangenome composition and unequal within-group heterogeneity. The gene composition of swine isolates exhibited the widest spread, indicating highest heterogeneity and diversity in the pangenome, while the composition of accessory genes of wild bird isolates was most homogenous, evidenced by the tightest clustering of the genomes (**Fig. 1d**). Overall, these results show that *S.* Typhimurium exhibits substantial variation in pangenome structure across isolation sources, with swine isolates displaying the most dynamic pangenome.

### 3.2 Positive selection in the core genome differed significantly among isolation sources in *S.* Typhimurium, but not in accessory genes

Different environments impose different selective pressures (e.g., receptors, chemical sanitizer, microbiota, temperature/pH/nutrients), which drive genomic adaptations and push pathogen populations toward niche specialization^76–80^. To quantify how positive selection shapes pangenome evolution of *S.* Typhimurium, we scanned core and accessory genes for selection and compared selection profiles across isolation sources. We found that positive selection was exerted on both core and accessory genes and was generally more common in accessory genes than core genes across sources, with wild birds as the exception (**Fig. 2a**). The proportion of accessory genes under positive selection was 0.82% among poultry isolates, while that of core genes was 0.78%, significantly higher than wild bird (0.55%), bovine (0.33%), and swine (0.22%) isolates (Fisher’s exact *P* = 0.0014; **Fig. 2a**). In comparison, the proportion of accessory genes under positive selection was not significantly different across isolation sources (*P* = 0.1765), ranging from 0.38% for swine isolates to 0.82% for poultry isolates (**Fig. 2a**). When all genes (core and accessory genes) were considered, the pattern mirrored that for core genes: the proportions of all genes under positive selection were significantly different across isolation sources (*P* = 0.0005); poultry isolates showed the highest proportion under positive selection (0.80%), followed by wild bird (0.52%), bovine (0.44%), and swine (0.28%) isolates (**Fig. 2a**). These results indicate that, although positive selection is more widespread in accessory genes in general, positive selection targeting core genes may play a more important role in source-specific adaptation in *S.* Typhimurium.

**Figure 2:**
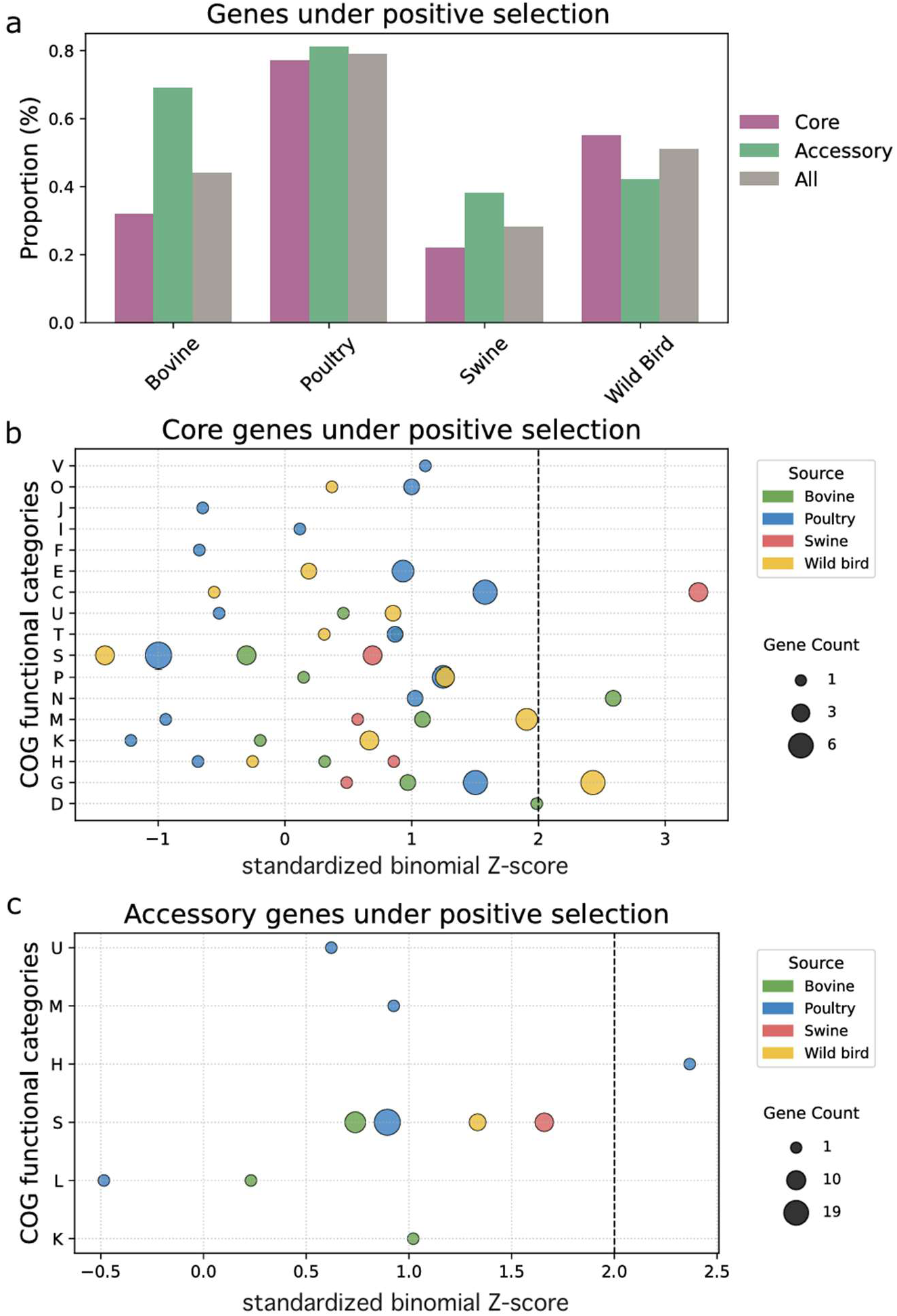
Degree and functional targets of positive selection in *S*. Typhimurium. (a) The proportion of core genes (purple), accessory genes (green), and total genes (grey) under positive selection in S. Typhimurium. Enrichment of COGs among (b) core genes and (c) accessory genes under positive selection. An standardized binomial Z-score greater than 2 (dashed line) indicates significant overrepresentation (*P* < 0.05). Circle size is proportional to the number of genes annotated to each COG. COG abbreviations are as follows: C: Energy production and conversion; D: Cell cycle control, cell division, chromosome partitioning; E: Amino acid transport and metabolism; F: Nucleotide transport and metabolism; G: Carbohydrate transport and metabolism; H: Coenzyme transport and metabolism; I: Lipid transport and metabolism; J: Translation, ribosomal structure, and biogenesis; K: Transcription; L: Replication, recombination, and repair; M: Cell wall/membrane/envelope biogenesis; N: Cell motility; O: Posttranslational modification, protein turnover, chaperones; P: Inorganic ion transport and metabolism; Q: Secondary metabolites biosynthesis, transport, and catabolism; T: Signal transduction mechanisms; U: Intracellular trafficking, secretion, and vesicular transport; V: Defense mechanisms.

To understand which gene functions tend to be subject to positive selection in *S.* Typhimurium and how they differ by isolation sources, we performed a binomial test to identify COGs significantly enriched among core and accessory genes under positive selection for a given source. Three COG categories were significantly enriched among core genes showing evidence of positive selection (*P* < 0.05; **Fig. 2b**). Within swine isolates, genes involved in energy production and conversion (C) were significantly enriched, whereas cell motility (N) was enriched in bovine isolates and carbohydrate transport and metabolism (G) in wild-bird isolates. Compared to core genes, only poultry isolates had one COG, coenzyme transport and metabolism (H), significantly enriched among accessory genes showing evidence of positive selection (**Fig. 2c**). The source-specific targets of positive selection on core gene functions in *S.* Typhimurium suggest adaptation to varied pressures it encounters in different conditions.

### 3.3 *S.* Typhimurium in swine harbors especially high burdens of MGEs and ARGs

MGEs are key drivers of pangenome plasticity and host adaptation in bacterial pathogens ^78–80^. To understand the associations between MGEs in *S.* Typhimurium and isolation source, we profiled prophages and plasmids in each genome and compared their prevalence across isolation sources. The proportions of genomes comprised of prophages, total and intact, were significantly different across sources (Kruskal–Wallis *P* = 1.27e-48 and 3.07e-49; **Fig. 3a** and **Fig. S2**, respectively). Swine isolates contained the highest proportions with total and intact prophages, followed by wild bird and bovine isolate genomes. Poultry isolates contained a significantly lower proportion of both total and intact prophages than isolates from other sources. Like prophages, the proportion of the genome comprised of plasmids was also significantly different across sources (Kruskal–Wallis *P* = 7.59e-47; **Fig. 3b**). Predicted plasmids were more prevalent among poultry and swine isolates on average than in isolates from other sources. With MGE burden acting as proxy for predicted HGT events, these results could indicate that *S.* Typhimurium experiences variable potential for genetic mobility in different environments, with the high prevalence of MGEs in swine isolates potentially contributing to its more open and larger pangenome that was discovered in this study.

**Figure 3:**
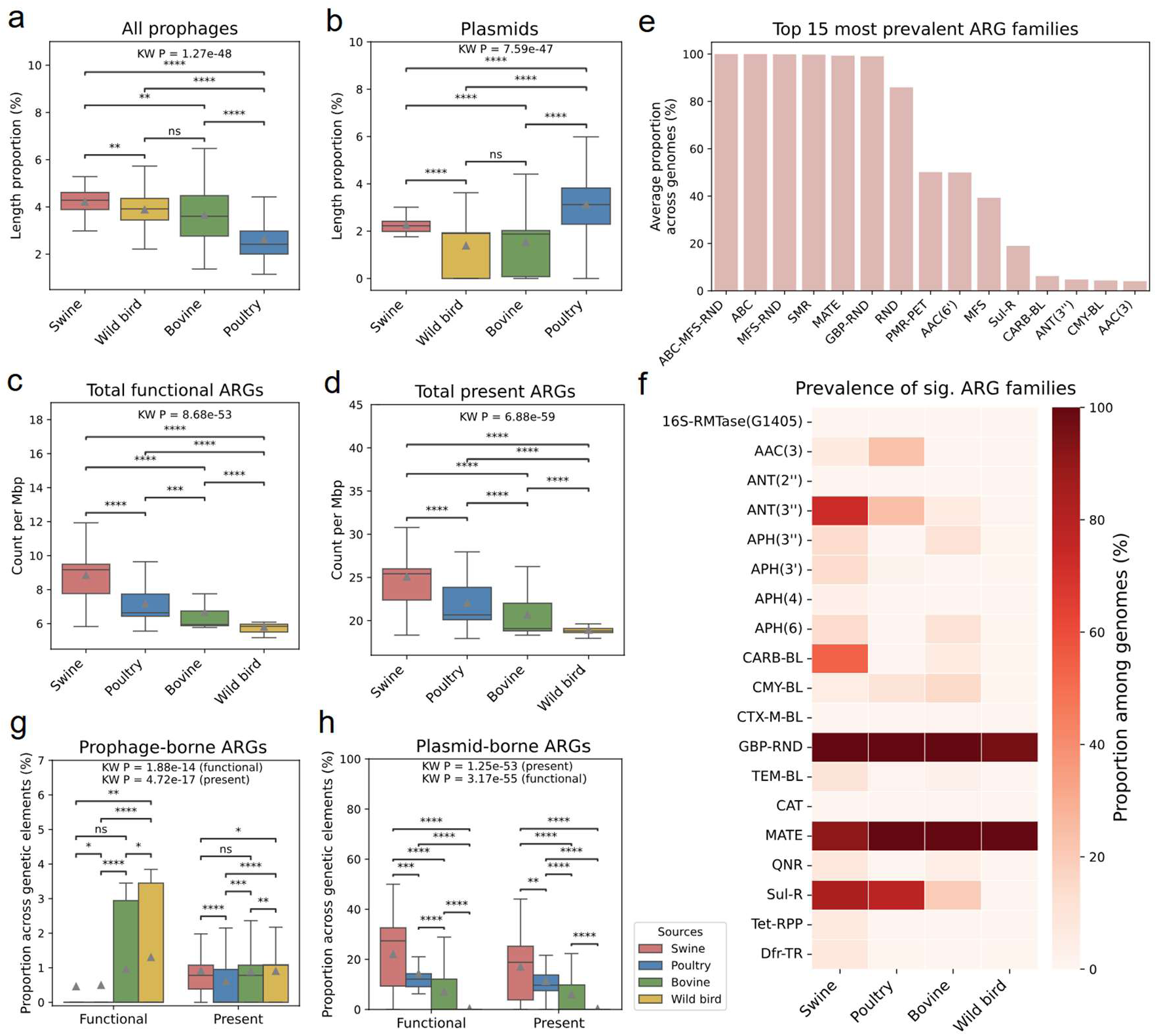
MGE and ARG profiles of *S*. Typhimurium. Prevalence of (a) prophages, (b) plasmids, and (c) functional ARGs, and (d) present ARGs compared across isolation sources. (e) The average proportion of the top 15 most prevalent ARG families among analyzed genomes. (f) The prevalence of ARG families significantly associated with isolation source. Significance was determined by Fischer’s exact tests comparing the prevalence of a given ARG family across isolation sources. A darker hue indicates a higher proportion of genomes containing that specific ARG family given an isolation source. The prevalence of functional and present ARGs predicted to originate from (g) prophages and (h) plasmids compared across isolation sources. For (a), (b), (c), (d), (g) and (h), box plots display the interquartile range (IQR) with the median indicated as a line and the mean indicated as a triangle and whiskers extending to 1.5 times the IQR. KW refers to the Kruskal-Wallis test. Significance levels are denoted by “*”, “**”, “***”, “****”, and “ns” for adjusted *P* < 0.05, < 0.01, < 0.001, < 0.0001, and ≥0.05 (not significant) for two-sided Mann-Whitney *U* tests for pairwise comparisons.

Rising AMR among *Salmonella* isolates poses a growing threat^81–83^. To assess the associations between AMR potential in *S.* Typhimurium and isolation source, we profiled ARGs and compared their prevalence across isolation sources. Significant differences were observed in both total count of putatively functional and present ARGs across isolation sources (*P* = 8.68e-53 and 6.88e-59; **Fig. 3c and 3d**, respectively). Swine isolates exhibited the highest number of ARGs, averaging a total of 26 present and 10 putatively functional ARGs per genome length (Mbp), followed by poultry and bovine isolates. Wild bird isolates contained the least ARGs, with less than 20 present and 6 functional ARGs per Mbp. Unique present and functional ARGs followed a similar trend, with swine containing the largest number, averaging slightly less than 18 and 8 ARGs per Mbp respectively, followed by poultry, bovine, and wild bird isolates being the lowest (**Fig. S3a** and **S3b**). These results are supported by the CARD-RGI analysis, in which ARGs were classified into three categories, “Perfect”, “Strict”, and “Loose”. Results show that the “Perfect” and “Strict” categories combined (**Fig. S4a and b**) correspond closely to the genes we classified as “putatively functional ARGs” (**Fig. 3c and Fig. S3b**). This result aligns with the definitions of the “Perfect” and “Strict” categories used in CARD-RGI, which indicate sufficient sequence support to encode a functional protein, and is therefore consistent with the definition of the “functional” category used in this study. Combining the “Perfect,” “Strict,” and “Loose” categories, consistent trend in the difference of ARGs among isolation sources was also observed (**Fig. S4c and d**) compared to our “present” category (**Fig. 3d and Fig. S3a**).

To identify AMR traits that are significantly different across isolation sources, we conducted Fisher’s exact tests on putatively functional ARGs and their families, drug classes, and resistance mechanisms. The prevalence of 47 ARGs was found to be significantly associated with isolation sources (adjusted *P* < 0.05 for all; **Fig. S3c**). Notably, 30 ARGs, such as *ANT(3’’)-lla*, *CARB-3*, *CARB-53*, *aadA2*, *aadA3*, cmlA9, *floR*, *qacE*, qacEdelta1, *sul1*, and *tet(G)*, were significantly more prevalent among swine isolates than other sources. For ARG families, four were detected in 100% of the isolates, such as ATP-binding cassette, major facilitator superfamily, and resistance-nodulation-cell division antibiotic efflux pumps (**Fig. 3e**). The prevalence of 19 ARG families was significantly associated with isolation sources (adjusted *P* < 0.05 for all; **Fig. 3f**). Swine isolates exhibited the most diverse ARG family profiles, with 12 being significantly more prevalent than other sources, such as ANT(3’’), APH(3’’), APH(3’), APH(6), CARB-BL, TEM-BL, and Tet-RPP (**Fig. 3f**). For drug classes, nine were detected in 100% of isolates, such as nitroimidazole and phosphonic acid antibiotics (**Fig. S5a, Table S2**). A total of 12 drug classes were significantly associated with isolation sources (adjusted *P* < 0.05 for all; **Fig. S5b, Table S2**). Consistent with ARG families, swine isolates exhibited the most diverse drug class profiles, with 5 being significantly more prevalent than other sources, including cephalosporin-monobactam-penam-penem (CMPP), diaminopyrimidine antibiotic (DAP), disinfecting agents and antiseptics (DAA), penam (PEN), phenicol antibiotic (PHN), and sulfonamide and sulfone antibiotics (SUL) (**Fig. S5b**). Lastly, for resistance mechanisms, there were seven different mechanisms identified across all 595 genomes, with ARGs involved in efflux and permeability encountered in nearly 100% of isolates and efflux alone in approximately 60% (**Fig. S5c**). Based on Kruskal–Wallis tests, count of ARGs belonging to six resistance mechanisms were significantly different among isolation sources (adjusted *P* < 0.05 for all; **Fig. S5d**). Swine isolates harbored ARGs with the most varied resistance mechanisms, with 30 ARGs per genome on average functioning via antibiotic efflux and nearly 11 via inactivation (**Fig. S5d**). These results indicate that ARGs in *S.* Typhimurium are consistent with possible antimicrobial selection in food-animal associated settings particularly swine, possibly attributed to the diverse antibiotics used.

To evaluate ARG mobility in *S.* Typhimurium, we determined whether any ARGs were likely carried on prophages and plasmids. A relatively small proportion of ARGs (less than 4% in general), both present and functional, were predicted to be present on prophages, with a significant difference among isolation sources (Kruskal-Wallis *P* = 4.72e-17 and 1.88e-14, respectively; **Fig. 3g**). Swine isolates had the highest and lowest average proportion of ARGs located on prophages for present and functional ARGs, respectively (**Fig. 3g**). *MdtK* was identified as a predominant functional ARG on prophages, comprising more than 80% among all functional ARGs located on prophages **(Fig. S6b**) and more than 20% among all functional *MdtK*s detected (**Fig. S6a)**. No functional ARGs were exclusively associated with prophages, and over 75% exhibited low prophage prevalence, accounting for less than 5% of all detected genes for a given ARG **(Fig. S6c)**. Compared to prophages, a much higher proportion of ARGs were predicted to be plasmid-borne, and it was significantly different among isolation sources as well (*P* = 11.25e053 and 3.17e-55 for present and functional ARGs, respectively; **Fig. 3h)**. Swine isolates contained the highest average predicted proportion of plasmid-borne ARGs (∼20% and 30% for present and functional ARGs, respectively). Among all predicted plasmid-borne functional ARGs, *tet(A)* and *sul2* were the most prevalent, with relative abundances of approximately 20% and 18%, respectively **(Fig. S7a)**. Unlike ARGs encoded within prophage regions, a large proportion (∼55%) of functional ARGs were exclusively identified on plasmids (**Fig. S7b)**. These results suggest that HGT, particularly through plasmids, may play a key role in ARG dissemination in *S.* Typhimurium, and the elevated ARG burden in swine isolates reflects frequent transfer via plasmids and past transduction events.

To determine if imbalanced source group size may cause bias in the associations between MGE and AMR profiles and isolation sources, we repeated the key MGE and ARG analyses using subsampling with N = 56 for each isolate source, corresponding to the number of swine isolates. Upon comparison to the complete data set (**Fig. 3**), we observed consistent relationships with isolation sources in the frequency of prophages (**Fig. S8a**), plasmids (**Fig. S8b**), functional ARGs (**Fig. S8c**), present ARGs (**Fig. S8d**), top 15 most prevalence ARG families (**Fig. S8e**), drug classes (**Fig. S8f**), and resistance mechanisms (**Fig. S8g**), and the frequency of prophage-borne (**Fig. S8h**) and plasmid-borne ARGs (**Fig. S8i**). In addition, to assess the potential bias imposed by temporal variability in the collection years, we also repeated these key MGE and ARG analyses using genomes with documented collection year of 2021. Notably, none of the wild-bird isolates were isolated from samples collected in 2021 (**Table S1**). Thus, only swine, bovine, and poultry isolates were included in these analyses. Upon comparison to the complete data set (**Fig. 3**), we observed consistent relationships with isolation sources in the frequency of prophages (**Fig. S9a**), plasmids (**Fig. S9b**), functional ARGs (**Fig. S9c**), present ARGs (**Fig. S9d**), top 15 most prevalence ARG families (**Fig. S9e**), drug classes (**Fig. S9f**), and resistance mechanisms (**Fig. S9g**), and the frequency of prophage-borne (**Fig. S9h**) and plasmid-borne ARGs (**Fig. S9i**). Overall, these results indicate that our main findings on MGEs and ARGs are robust to sampling imbalance across isolate source categories and temporal variability in the collection years.

### 3.4 Machine learning models identify genetic variants predictive of isolation source of *S.* Typhimurium

As we observed strong associations between genome content and isolation source in *S.* Typhimurium, we hypothesized that machine learning models could be utilized to identify genetic variance predictive of isolate source. To test this hypothesis, we compared several machine learning algorithms, such as decision tree, random forest, lightGBM, and support vector machine (see **Methods**). After hyperparameter tuning, a lightGBM model was identified as the best model for predicting isolation sources of genomes with gene presence/absence as features (**Table S3**). This model trained 50 boosted trees with a learning rate of 0.1, which were constrained by 3 leaves and a depth of 10. Generalization was helped by L2 regularization and a minimum split gain of 0.2, so weak splits were pruned. Sampling-wise, it used row subsampling at 0.9, and all features per tree. This model achieved a mean auROC of 0.97, 1.00, 0.98, and 1.00 (**Fig. 4a**) and a mean auPR of 0.80, 1.00, 0.90, and 1.00 (**Fig. 4b**) for predicting bovine, poultry, swine, and wild bird origins using the test set, respectively. To interpret outputs from this best-performing model, we employed SHAP to evaluate the importance of each feature in prediction. The top ten most predictive features for all isolation sources included accessory genes *ydcX* and *cusB*, encoding a GhoT/OrtT-family membrane-associated toxin and a periplasmic adaptor protein of the CusCBA copper/silver efflux system, respectively (**Fig. 4c**, **Table S5**). Also, we found that genes most predictive of each source were different (**Figs. S10a – S10d, Table S5**). The top three predictive genes for each isolation source included *cusB* for bovine, *gatB* (encoding a PTS galactitol-specific IIB component involved in carbohydrate transport and metabolism) for poultry, *ramB* (encoding sequence-specific DNA-binding transcriptional regulator) for swine, and *ydcX* and *hdeB* (encoding acid-stress chaperone) for wild bird. Several accessory genes identified here, including *cusB*, *ydcX*, and *hdeB* are associated with stress response.

**Figure 4:**
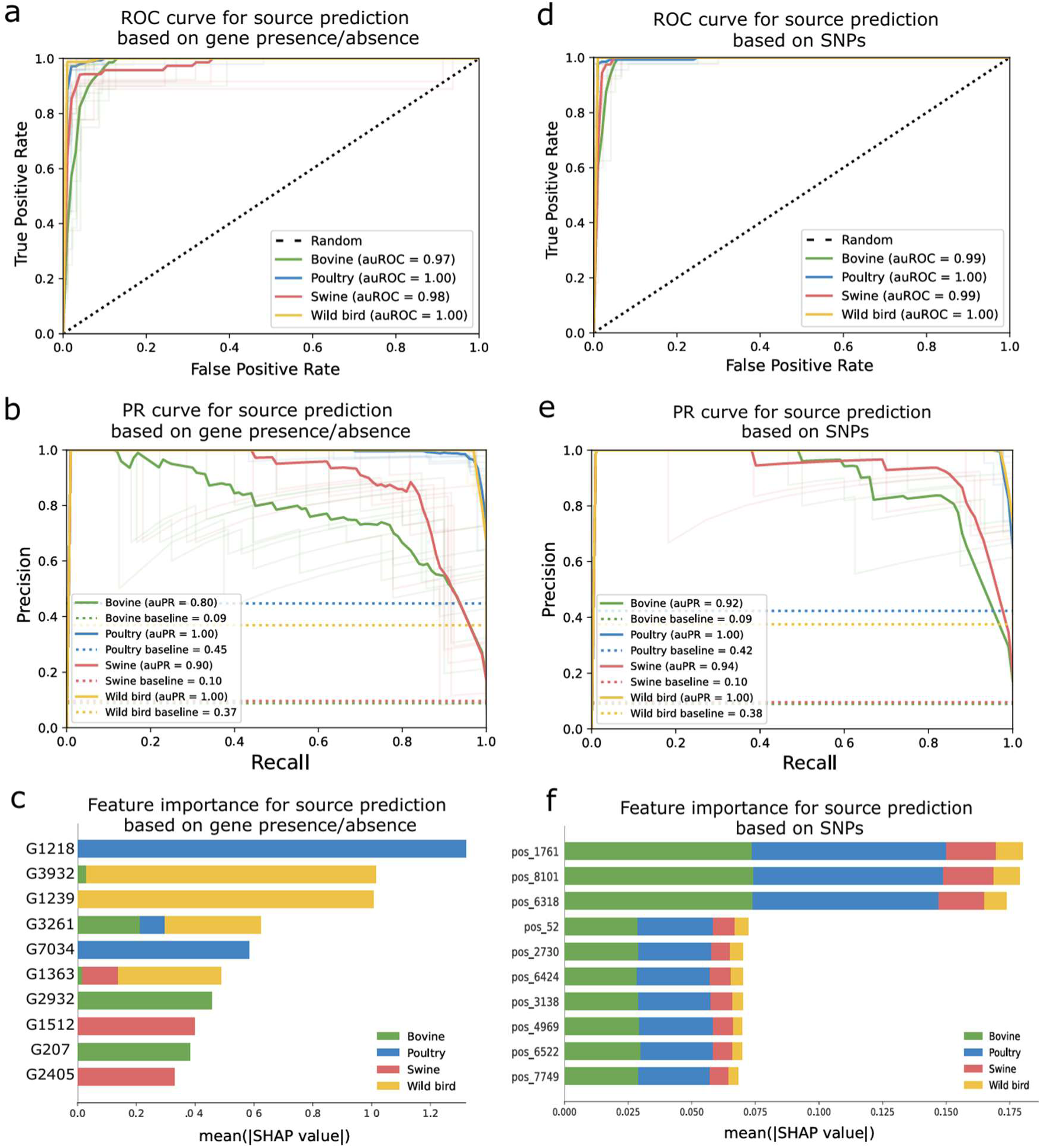
Machine learning models trained on gene presence/absence and core SNP profiles of *S.* Typhimurium for isolation source prediction. (a) Receiver operating characteristic (ROC) curves and (b) precision-recall (PR) curves for the prediction of isolation sources from gene presence/absence profiles using a lightGBM model. (c) The top ten most predictive genes for all isolation sources (SHAP-based, X-axis), sorted by descending importance. (d) Receiver operating characteristic (ROC) curves and (e) precision-recall (PR) curves for the prediction of isolation sources from core SNPs using a SVM model. (f) The top ten most predictive SNP locations for all isolation sources (SHAP-based, X-axis), sorted by descending importance. For (a), (b), (d), and (e), each curve reflects one evaluation using test data, repeated 10 times, denoted by light color lines. The dark color line on the ROC curve denotes random chance accuracy. The colored dotted lines on the PR curves denotes class prevalence baselines. For (c) and (f), the functional annotations of the predictive genes and SNPs are provided in **Table S5** and **S6**, respectively.

In addition to using gene presence/absence as features, we also developed a machine learning model using core SNPs as features and identified SVM as the best model after hyperparameter tuning (**Table S4**). This model utilizes a polynomial kernel with degree 4, a regularization parameter of 0.5, and the shrinking optimization option enabled. This model achieved a mean auROC of 0.99, 1.00, 0.99, and 1.00 and a mean auPR of 0.92, 1.0, 0.94, and 1.00 for predicting bovine, poultry, swine, and wild bird origins using the test set, respectively (**Figs. 4d and 4e**). Based on SHAP, the top three most predictive features for all isolation sources included SNPs Pos1761, Pos8101, and Pos6318, which are located on core genes *kdpD* (encoding a sensor kinase involved in potassium homeostasis and osmotic stress response), *adiY* (encoding an AraC/XylS-family transcriptional regulator involved in acid resistance), and *relA* (encoding a (p)ppGpp synthetase that regulates the bacterial stringent response to nutrient limitation and other environmental stresses), respectively (**Fig. 4f**; **Table S6**). The core SNPs most predictive of isolation source shared some similarities among bovine, poultry, and swine origins (**Figs. S11a – S11d, Table S6**). Core genes where the top three predictive SNPs were located for each isolation source included *relA* and *adiY* for both bovine and poultry, *adiY* and *dsbG*, involved in disulfide bond formation for swine, and *ung*, which encodes a uracil-excising protein*, yhjC*, which is involved in LysR substrate binding, and *cbiQ,* which encodes a cobalt transfer protein for wild bird. Several genes identified here, including *kdpD*, *adiY*, *relA*, and *dsbG* are associated with stress response. Overall, our results suggest that both gene presence/absence and core SNP patterns, particularly those involved in stress response, could enable strong source discrimination in *S.* Typhimurium using machine learning.

To ensure that the machine learning models are not simply learning sampling structure, we externally validated the best models based on gene presence/absence and core SNPs using isolates collected from 2002-2020. Results show that both models remained robust in distinguishing between isolation sources (auROC > 0.9 for all; **Fig. S12a-b**). However, compared to the performance of the test set (**Fig. 4b and d**), while auPR values remain high for most source groups, it decreased noticeably for the swine classes (**Fig. S12c-d**), suggesting reduced precision-recall performance for this source. This reduction may be partly attributable to smaller sample size in the validation set. Overall, source discrimination remains strong, as reflected by the overall high auROC values and auPR values, supporting the robustness of our models.

## 4 Discussion

Many bacteria species exhibit substantial genomic plasticity, with gene gain and loss at multiple scales facilitating adaptation to stressors encountered in diverse environments or hosts^84^. To understand how genomic diversification may contribute to differential adaptation in foodborne pathogens, we analyzed 595 genomes of *S.* Typhimurium representing four major sources, including three food animal sources - bovine, poultry, and swine, and one non-food animal source - wild birds. We show that gain and loss of accessory genes, including ARGs, movement of which is associated with MGEs, particularly plasmids, together with positive selection acting on core genes, may jointly contribute to the differential adaptation of *S.* Typhimurium to diverse ecological niches.

We show that food animal isolates (i.e., swine, poultry, and bovine) generally exhibited a more open pangenome and higher ARG prevalence than the non-food animal source (i.e., wild birds). The elevated ARG prevalence is consistent with sustained antibiotic selection pressure within food production systems^85–87^. While prophylactically administering antibiotics broadly or including them in livestock feed to supplement growth was banned within the United States in 2017 by the Food and Drug Administration, the effects of these practices linger throughout the country^88^. Numerous diverse antibiotics have been implemented in various agricultural settings, with some, such as tetracycline and sulfonamides, being considerably more popular, and therefore, seeing a more dramatic decrease in their effectiveness, an issue of magnitude given their nature as broad-spectrum antibiotics^89,90^. According to previous studies, approximately 80% of the antibiotics sold in the United States are devoted to livestock or farming^90–92^. In addition to leaving trace amounts in food products that later are consumed by humans^90^, roughly 75% of any antibiotic used in livestock feed are not absorbed but rather excreted, contaminating the environment by infesting the soil, the water, and even other nearby plants with antibiotics, further contributing to elevated selective pressure and increasing rates of resistance^93^. Additionally, though increasingly regulated by the United States and by many countries, the continuing practice of spraying or otherwise utilizing antibiotics such as streptomycin in farming likewise contributes to the higher selective pressure faced by bacteria in agricultural environments^94,95^.

Notably, *S.* Typhimurium swine isolates exhibited the highest ARG prevalence in this study, suggesting decreased susceptibility, pending confirmation by phenotypic testing. This pattern can be attributed to a variety of farming practices, most of which revolve around the utilization of antimicrobials and antibiotics for growth and treatment and prevention of disease in swine production^96,97^. Studies have shown that, in the past, up to 88% of swine received antibiotics in their feed, intended to prevent disease and promote growth^90,91^. Many of the antibiotics historically utilized in swine farming – such as tetracyclines, amoxicillin, penicillin, streptomycin, and sulfamethazine – were present in nearly 50% or more pig feeds in some cases ^98–100^, likely explaining their elevated prevalence of associated ARGs in *S.* Typhimurium swine isolates observed in this study. Among them, streptomycin is not often prescribed for salmonellosis but has been used as a growth factor for swine farming^97^. Additionally, many of these drug classes correlate to the frequently seen *Salmonella* multi-drug resistant (MDR) strains from swine, including tetra- (ampicillin, streptomycin, sulfonamides, and tetracyclines (ASSuT)) and penta- (ampicillin, chloramphenicol, streptomycin, sulfonamides, and tetracyclines (ACSSuT)) resistant strains^97^. While chloramphenicol has been banned in pig feed in the United States, it is predicted that the introduction of the drug and subsequent linkage to other resistance – namely macrolides and aminoglycosides – could be contributing to the prevalence of chloramphenicol resistance in these MDR strains^101^. Further compounding this spread is the environmental conditions that feed back into the cycle. *Salmonella* can survive in the environment in waste matter for a prolonged period, during which the swine may ingest potentially antimicrobial-resistant *Salmonella* through the fecal-oral route of transmission^102^.

In addition to extrinsic factors such as antibiotic usage, we found that intrinsic factors, specifically plasmids-associated HGT, may play an essential role in the dissemination of ARGs among *S.* Typhimurium swine isolates. It has been reported that many ARGs are introduced by plasmid incorporation or, in the case of *Salmonella*, by conjugative or mobilizable plasmids^103,104^. This is consistent with our results, where nearly 40% of the ARGs on average detected across the swine isolates were predicted to be plasmid-borne, and a large proportion (∼55%) of putatively functional ARGs were exclusively detected on contigs predicted as plasmids. With *Salmonella*’s lack of competence when taking up free DNA, alternative mechanisms of plasmids-mediated transfer become the main avenue of AMR gene transmission^103^. Since the plasmids are sizable and require numerous resources, to survive, they may provide a clear evolutionary advantage, which ARGs do, when faced with antibiotics and the risk of susceptibility^105,106^.

In addition to plasmids, we also identified evidence that historical transduction may be important to the dissemination of ARGs among *S.* Typhimurium swine isolates. Prophages are commonly found in *S.* Typhimurium genomes, with approximately 99% being capable of generalized transduction^107,108^. Even though transduction may not be the main mechanism of HGT of ARGs in *S.* Typhimurium, it poses a significant health risk^107,109^. It has been reported that some *S.* Typhimurium strains exhibit a MDR penta-resistance to the ACSSuT phenotype, which is theorized to be transduced by two P22-like prophages^32,110^. Additionally, there is evidence that fluoroquinolones cause DNA damage to the bacteria, inducing an SOS response, which can increase gene expression of prophage and induce a prophage, potentially driving HGT of ARGs in *Salmonella* inadvertently^111^. In summary, Historical and ongoing HGT may contribute to the high prevalence of ARGs in *S.* Typhimurium within swine production systems.

In addition to ARG dissemination, MGEs likely contribute to the more open pangenome observed in swine isolates as well, which reflects broader evolutionary dynamics governing bacterial genome plasticity, where adaptation can occur through gene loss or gain. We hypothesize that this pattern could be attributed to combined effects of high host density, frequent microbial encounters, stable gut environments, and repeated ecological bottlenecks, which together favor horizontal acquisition and long-term retention of accessory genetic material^16,78^. Gene loss (or degradation) is typically considered to be an evolutionary driver, aimed at streamlining function and reducing redundancy within the bacteria^112^. This can be seen strongly in host-restricted pathogens, where, due to the constancy of the environment, the need to constantly adapt is diminished^113,114^. Contrary to that, gene gain, often seen in more generalist pathogens, is a measure of genome flexibility^115^. A more open pangenome indicates a greater capacity for adaptation through gene acquisition and diversification, possibly permitting strains to gain host-specific functions such as virulence factors, metabolic flexibility, and stress response mechanisms – needed to adapt to the diverse array of challenges presented across varying hosts^116^. While host adapted *Salmonella* serotypes are associated with genome degradation – including the formation of pseudogenes (non-expressed remnants of previous genes), there are lower gene loss rates in generalist *Salmonella* serovars^117^. Together, these dynamics suggest that acquisition of novel genes within an open pangenome may support adaptation to host-associated environments, potentially contributing to the elevated accessory genome content observed in swine isolates.

Positive selection drives bacterial adaptive evolution by favoring beneficial mutations that enhance fitness under specific environmental pressures, enabling rapid optimization of functions and improved survival across changing niches. In this study, accessory genomes exhibited a higher degree of positive selection than core genomes across most isolation sources. Given the open nature of accessory genomes and their frequent gain and loss driven by environmental exposure and functional necessity, this pattern is expected as newly acquired genes are often deleterious, and strong positive selection is required to retain only those variants that confer adaptive benefits^13,118,119^. In contrast, the core genome of *S.* Typhimurium was found to be highly conserved and primarily encoding essential cellular functions, making it less prone to frequent innovation and strongly shaped by purifying (or negative) selection to maintain established functionality^28,46,120^. Despite this overall trend, the frequency of core genes under positive selection differed significantly among isolation sources. This difference may reflect source-specific environmental conditions that expose *S. Typhimurium* to distinct combinations of temperature fluctuations, chemical pressures, and biotic interactions, thereby imposing different selective pressures on core functions across sources.

Genes and core SNPs predictive of the isolation source of *S.* Typhimurium were identified based on machine learning models, LightGBM and SVM, respectively. Many of the most predictive genomic features were associated with environmental stress adaptation, suggesting that source-specific ecological pressures may contribute to genomic differentiation in *S. Typhimurium*. Notably, highly predictive core SNPs occurred in *kdpD*, *adiY*, and *relA*, which participate in osmotic/potassium homeostasis, acid resistance, and the stringent response, respectively. The KdpD/KdpE system integrates potassium homeostasis with adaptation to osmotic and host-associated stresses, while AdiY regulates the arginine-dependent acid resistance pathway required for survival under low-pH conditions^121–123^. Acid adaptation is particularly relevant because *Salmonella* encounters acidic conditions across both host and environmental settings, and acid tolerance can influence survival under additional stresses^124^. Several source-predictive accessory genes were also associated with stress adaptation, including *cusB* for metal stress tolerance, *hdeB* for acid-stress protection, and *ydcX*, a GhoT/OrtT-family toxin linked to stress-associated growth modulation. Together, these patterns suggest complementary modes of source adaptation, with sequence-level selection modifying conserved core functions while accessory-gene gain and loss provide more flexible niche-specific capabilities. This interpretation is consistent with previous pangenome-wide analyses showing that both core-genome variants and accessory genes contribute to animal-source adaptation in *Salmonella*^125^. Since machine learning algorithms usually require a great deal of resources and time, identifying biomarkers with high discrimination power can reduce data input size, resulting in a quicker output^126^. Overall, these machine learning models and identified predictive genomic features could greatly benefit the food industry’s ability to track outbreaks and conduct source attribution in a timelier manner.

## 5 Conclusions

Here, we show evidence that *Salmonella* Typhimurium adapts to a broad range of ecological niches, spanning both food animal and non-food animal sources, through extensive genomic diversification. The elevated prevalence of antimicrobial resistance genes in food animal isolates, particularly those from swine production systems, may reflect selective pressures associated with antimicrobial use in animal production. We also suggest a central role for horizontal gene transfer mediated by plasmids in driving antimicrobial resistance dissemination as well as broader patterns of accessory gene gain and loss. Although core genomes were largely conserved in size across isolation sources and experienced less frequent positive selection than accessory genomes, selection on core genes may play a more important role in differential adaptation. These distinct evolutionary forces, including gain and loss of accessory genes and positive selection on core genes, likely enable adaptation of *S.* Typhimurium to diverse host and environmental contexts. Consistent with this notion, many genomic features highly predictive of isolation source identified by machine learning models are associated with environmental stress adaptation. These findings provide a foundation for understanding host-associated genomic patterns, but confirmation will require additional studies using complete assemblies and expanded sampling. Collectively, these findings advance our understanding of the genomic and evolutionary mechanisms shaping the broad ecological range of *S.* Typhimurium and demonstrate how genomic data can be harnessed to improve source tracking, enabling more efficient outbreak response in food production systems.

## Data availability

The genomic data and metadata used in this study were retrieved from the NCBI Pathogen Detection database. The accession numbers of genomes are listed in **Table S1**.

## Acknowledgments

We thank all members of the LEAPH laboratory for their enriching discussions. This work was funded by the Virginia Tech Engineering Faculty Organization-Opportunity (EFO-O) Seed Grant (JL, HL, AP, and RC), the Virginia Tech Global Change Center Undergraduate Research Grant (LO), and USDA FACT-CIN Award No. 2021-67021-34343 (SC and SL). The funders played no role in the study design, data collection, analysis, and interpretation of data, or the writing of this manuscript.

## Competing interest

All authors declare no financial or non-financial competing interests.

## Author contributions

JL designed the study. LO, SC, YG, and HZ analyzed the data with input from RC, SL, and JL. LO and JL wrote the paper with input from XD, AP, RC, and SL.

## Notes

### Competing Interest Statement

The authors have declared no competing interest.

### Summary of Updates

Revised in response to peer review comments. We refined the study design by imposing tighter QA/QC metrics and eliminating one source group due to high within-group heterogeneity. Additionally, we have included new analyses and expanded validation datasets to strengthen the core findings.

