## Supplemental Figures 1-12 for "The broad ecological range of *Salmonella enterica* serotype Typhimurium is associated with genomic diversification"

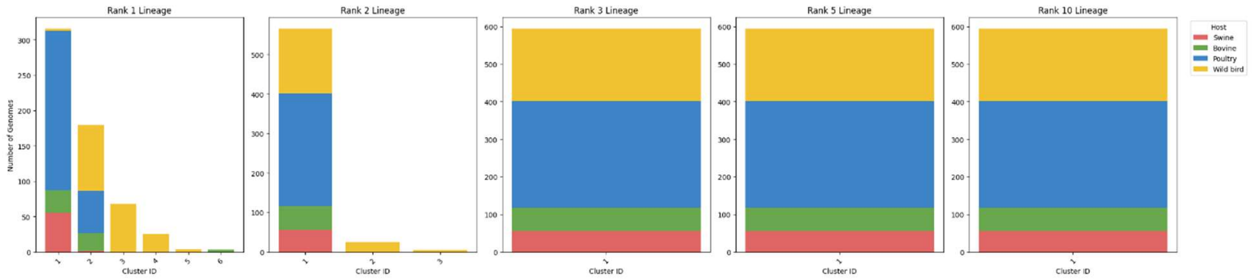


**Figure S1: Lineage clusters stratified by isolate source in *S.* Typhimurium.** Lineage clusters shown for *S.* Typhimurium isolates at Ranks 1, 2, 3, 5, and 10, stratified by host (swine – red, bovine – green, poultry – blue, wild bird – yellow). Rank refers to the resolution of the model, with Rank = 1 being the most specific.

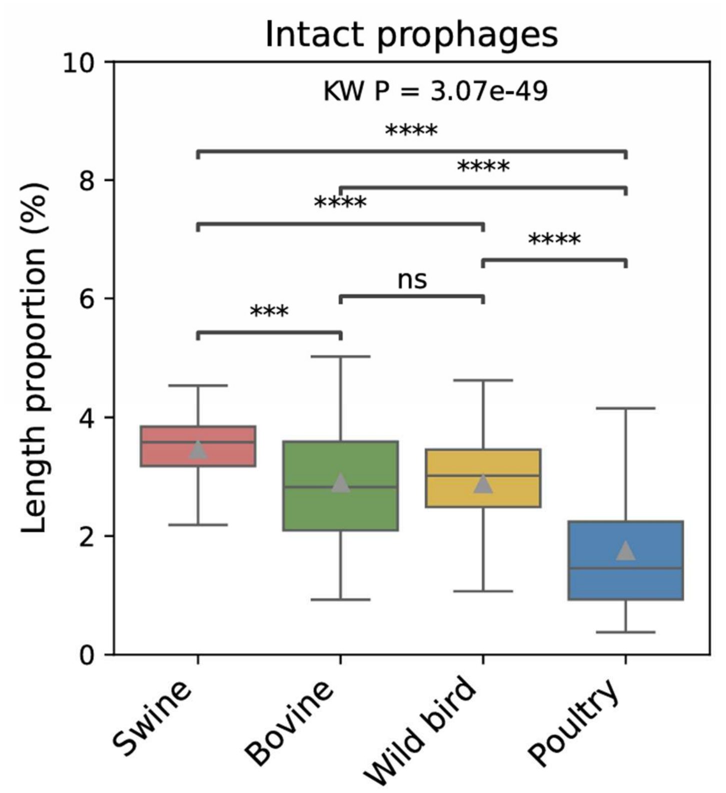


**Figure S2: Prevalence of intact prophages in *S*. Typhimurium compared across isolation sources.** Box plots display the interquartile range (IQR) with the median indicated as a line and the mean indicated as a triangle and whiskers extending to 1.5 times the IQR. KW refers to the Kruskal-Wallis test. Significance levels are denoted by "*", "**", "***", "****", and "ns" for adjusted $P < 0.05$, $< 0.01$, $< 0.001$, $< 0.0001$, and $\geq 0.05$ (not significant) for two-sided Mann-Whitney *U* tests for pairwise comparisons.

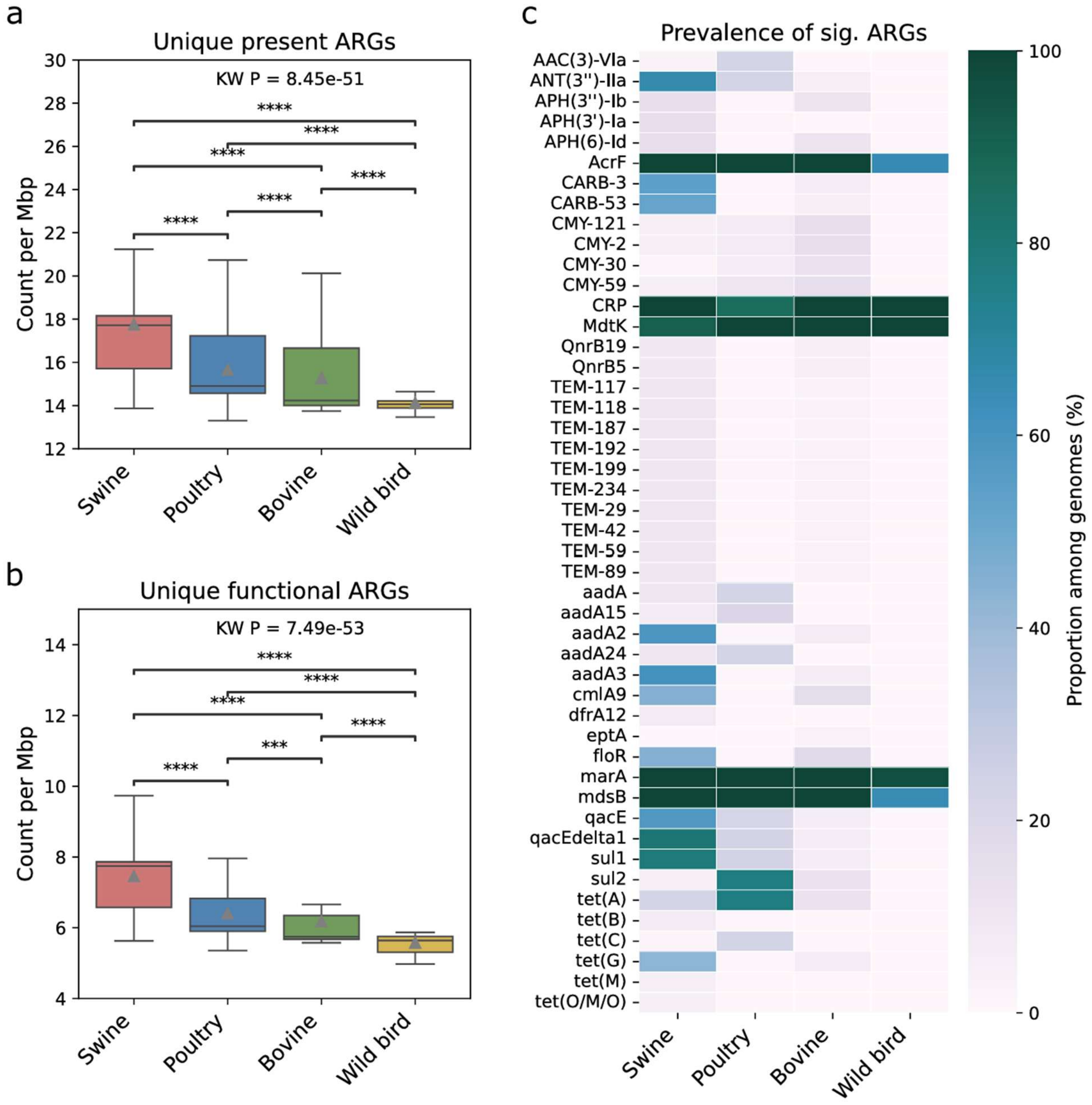


**Figure S3: Prevalence of ARGs in *S.* Typhimurium compared across isolation sources.** Prevalence of unique (a) present and (b) functional ARGs. (c) Prevalence of individual ARGs significantly associated with isolation source. Significance was determined by Fischer's exact tests comparing the prevalence of a given ARG across isolation sources. A darker hue indicates a higher proportion of genomes containing that specific ARG given an isolation source. For (a) and (b), box plots display the interquartile range (IQR) with the median indicated as a line and the mean indicated as a triangle and whiskers extending to 1.5 times the IQR. KW refers to the Kruskal-Wallis test. Significance levels are denoted by "*", "**", "***", "****", and "ns" for adjusted *P* < 0.05, < 0.01, < 0.001, < 0.0001, and ≥0.05 (not significant) for two-sided Mann-Whitney *U* tests for pairwise comparisons.

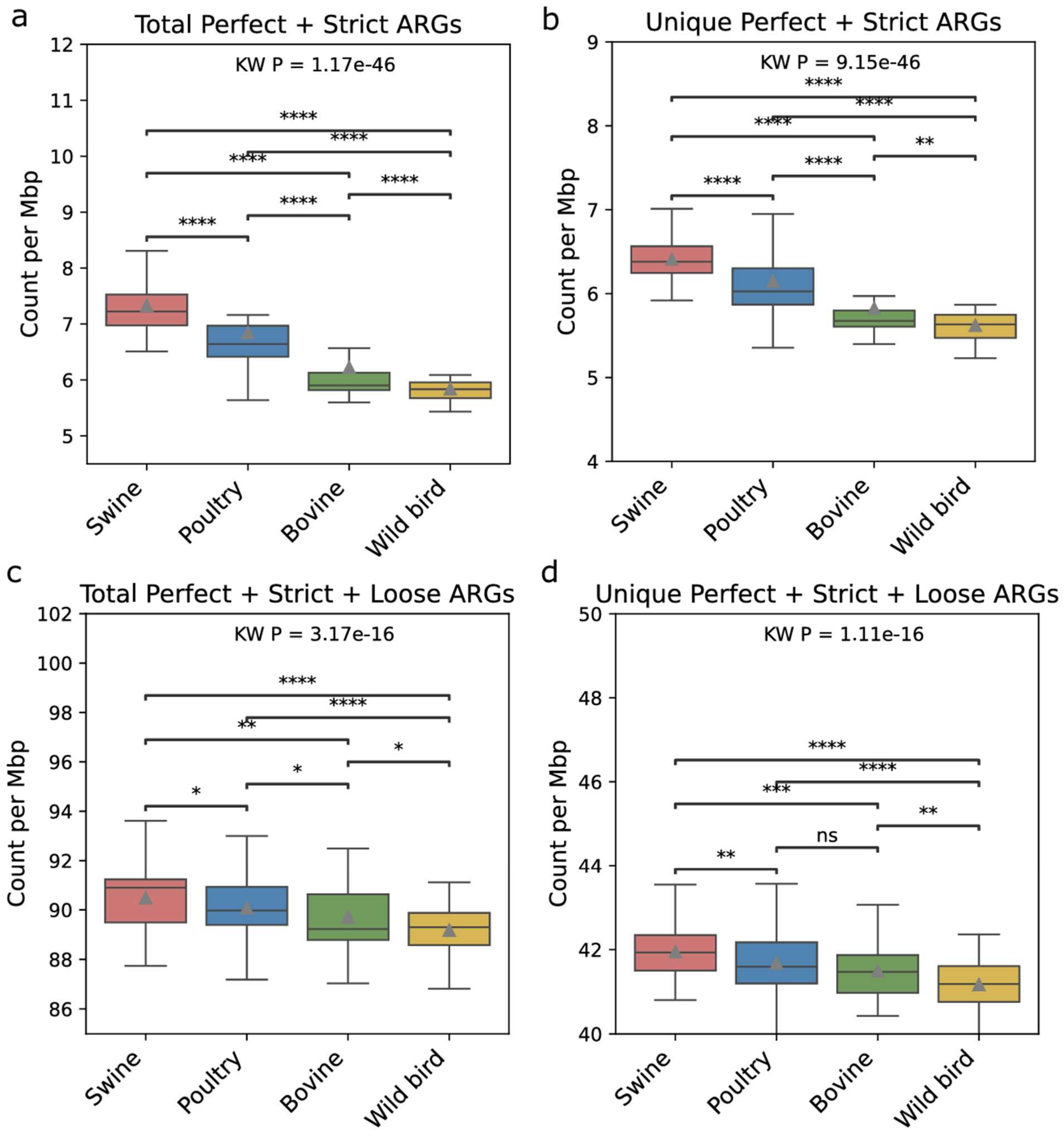


**Figure S4: CARD-RGI validation of AMR prediction pipeline.** (a) Total and (b) unique perfect and strict ARGs identified by CARD-RGI by host. (c) Total and (d) unique perfect, strict, and loose ARGs identified by CARD-RGI. Box plots display the interquartile range (IQR) with the median indicated as a line and the mean indicated as a triangle and whiskers extending to 1.5 times the IQR. KW refers to the Kruskal-Wallis test. Significance levels are denoted by "*", "**", "***", "****", and "ns" for adjusted $P < 0.05$, $< 0.01$, $< 0.001$, $< 0.0001$, and $\geq 0.05$ (not significant) for two-sided Mann-Whitney $U$ tests for pairwise comparisons.

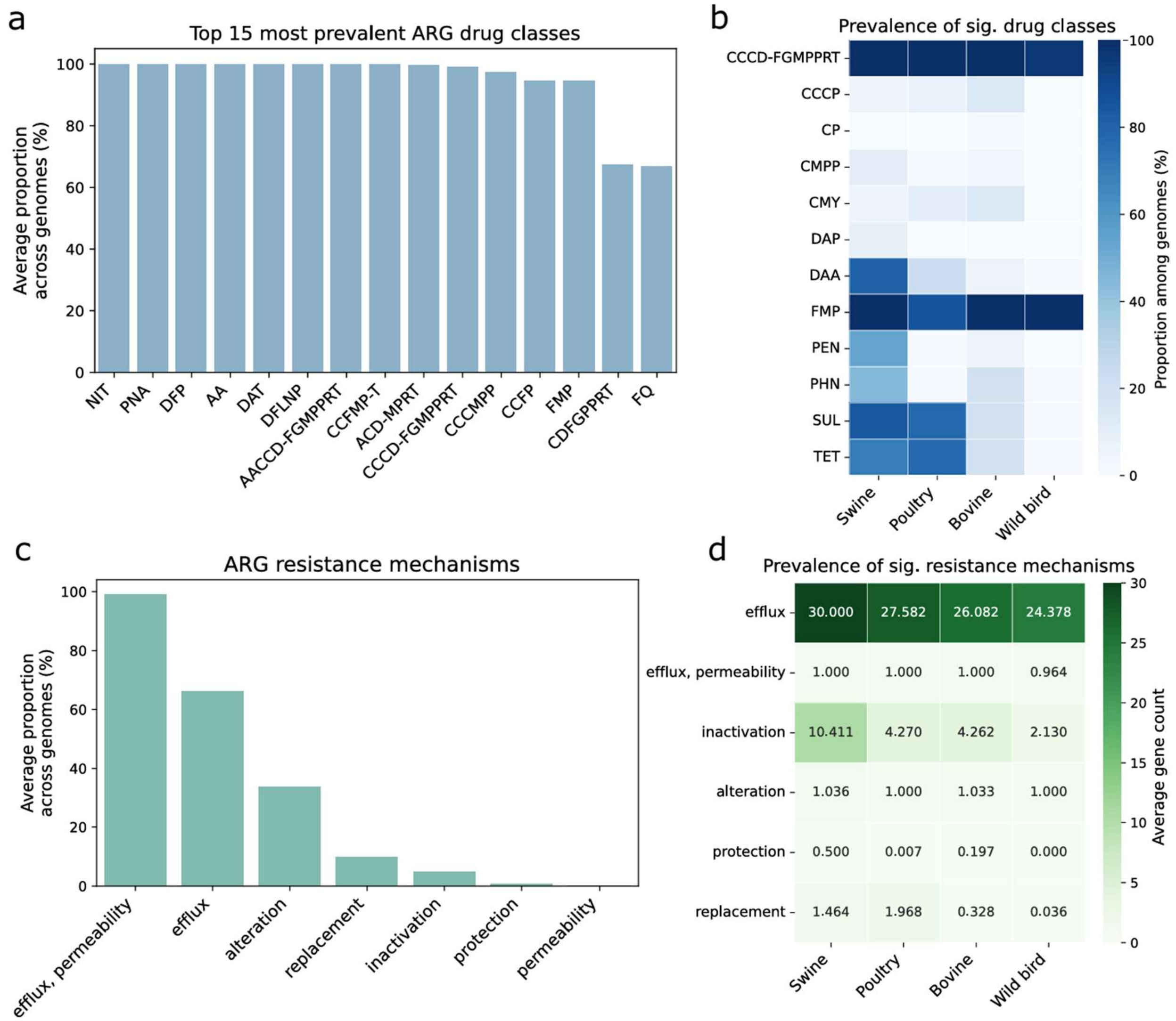


**Figure S5: Prevalence of ARG drug classes and resistance mechanisms in *S.* Typhimurium.** (a) The average proportion of the top 15 most prevalent ARG drug classes among analyzed genomes. (b) Prevalence of ARG drug classes significantly associated with isolation source. Significance was determined by Fischer's exact tests comparing the prevalence of a given ARG drug class across isolation sources. A darker hue indicates a higher proportion of genomes containing that specific ARG drug class given an isolation source. (c) Prevalence of ARG resistance mechanisms among analyzed genomes. The full names of the drug classes are provided in **Table S2**. (d) Prevalence of ARG resistance mechanisms significantly associated with isolation source. Significance was determined by two-sided Mann-Whitney *U* tests comparing the number of genes expressing a given resistance mechanism across isolation sources. A darker hue indicates a higher number of genes expressing that mechanism on average among genomes given an isolation source.

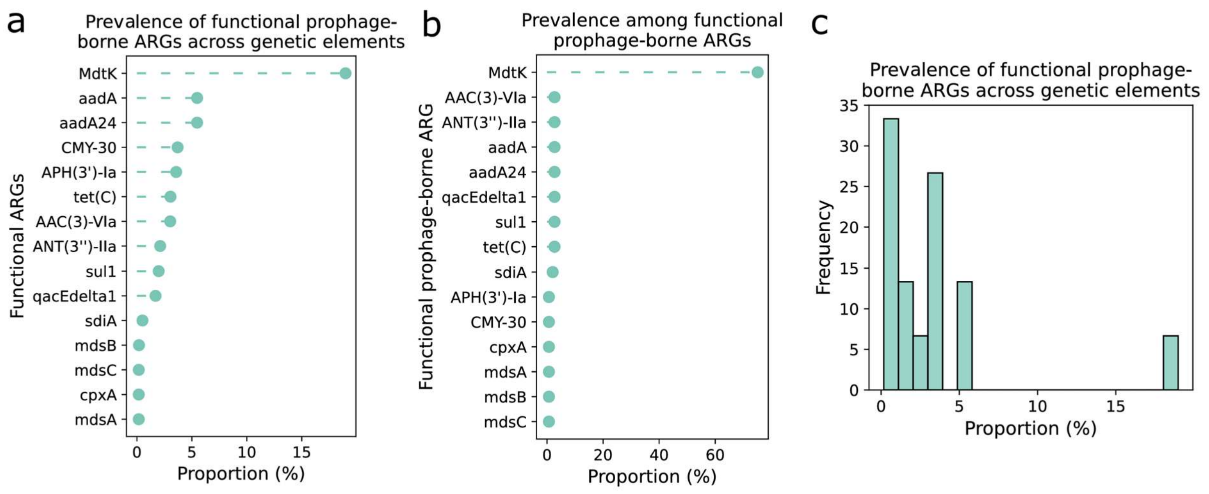


**Figure S6: Prevalence of functional, prophage-borne ARGs in *S*. Typhimurium.** Top 15 most prevalent prophage-borne ARGs (a) among all prophage-borne ARGs, and (b) across genetic elements harboring a given ARG, sorted by descending order. (c) Histogram showing the distribution of the prevalence of functional, prophage-borne ARGs compared across genetic elements harboring a given ARG.

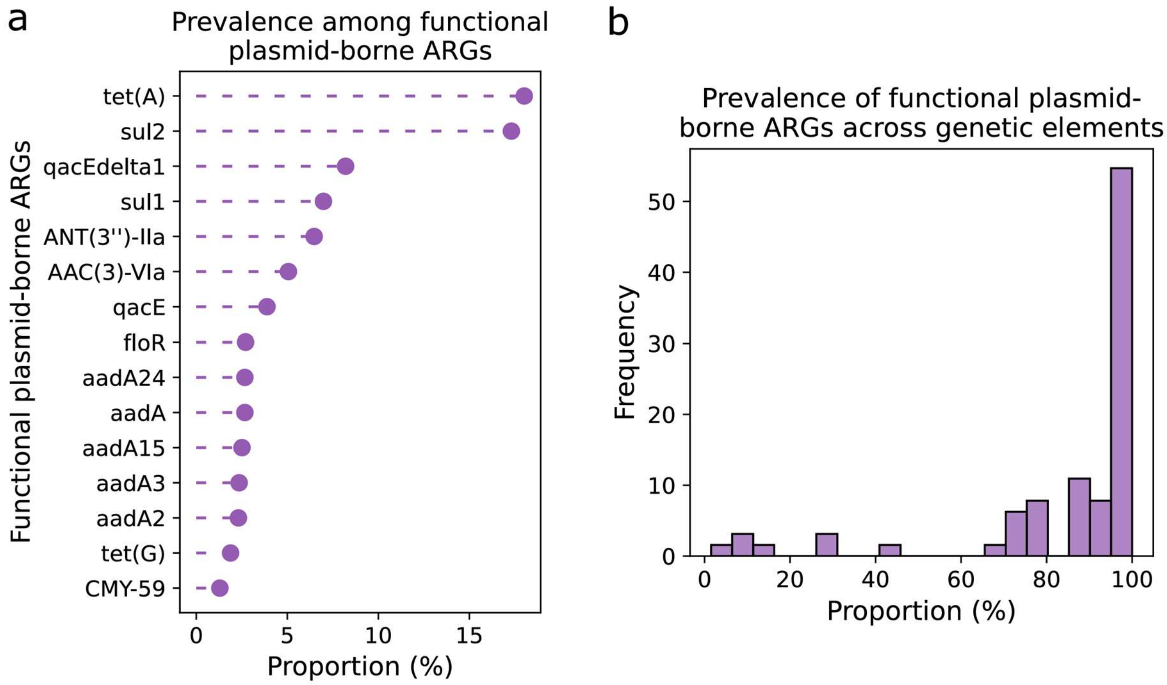


**Figure S7: Prevalence of functional, plasmid-borne ARGs in *S.* Typhimurium.** (a) Proportion of top 15 most prevalent plasmid-borne ARGs among all plasmid-borne ARGs. (b) Histogram showing the distribution of the prevalence of functional, plasmid-borne ARGs across genetic elements harboring a given ARG.

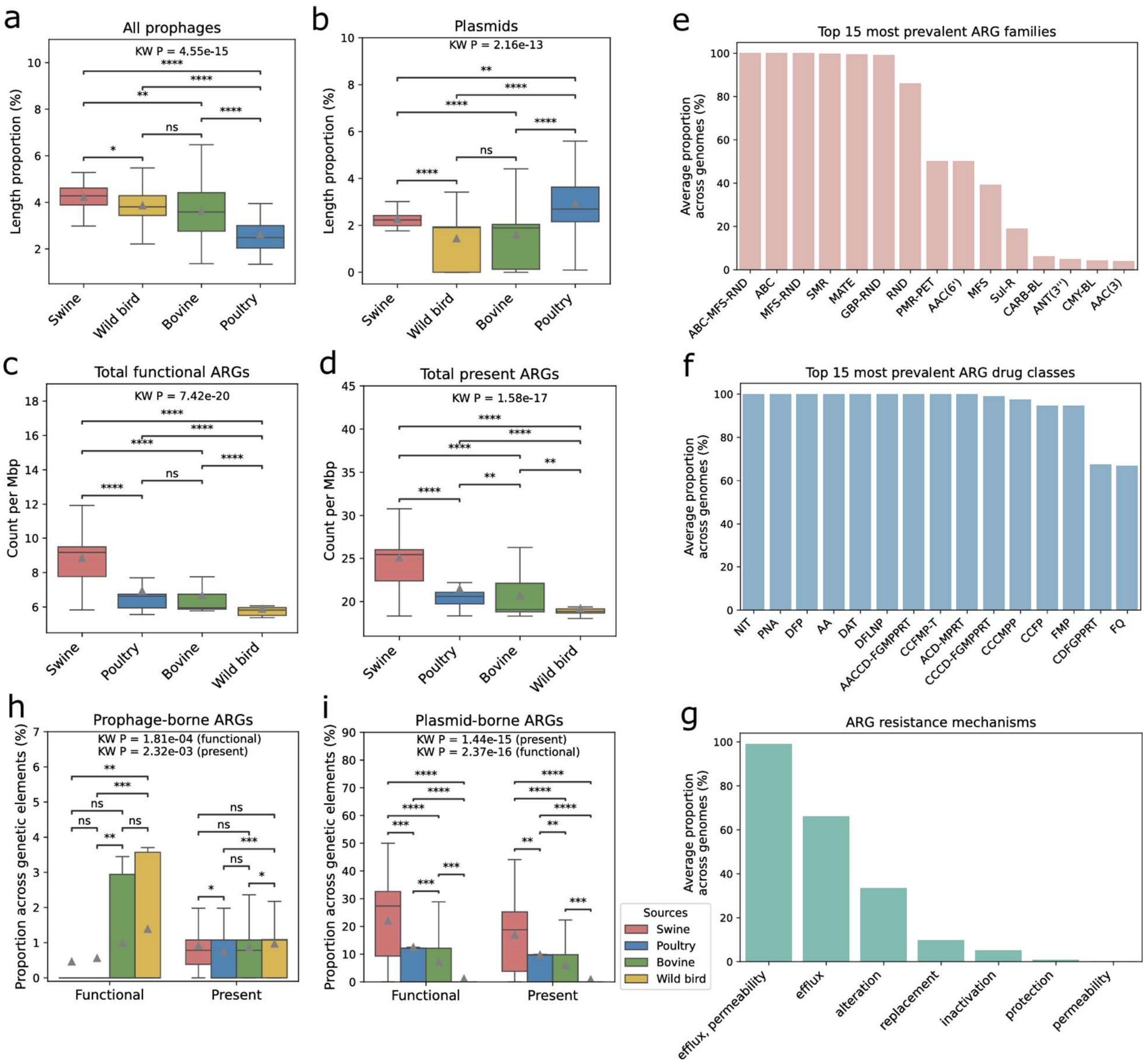


**Figure S8: AMR and MGE results for subsampled (N=56) *S.* Typhimurium genomes.** Length proportion of all identified (a) prophages and (b) plasmids identified on swine (red), wild bird (yellow), bovine (green), and poultry (blue) isolates. Total (c) putatively functional and (d) present ARGs per genome. Average proportion of the top 15 most prevalent (e) ARG families, (f) ARG drug classes, and (g) ARG resistant mechanisms. The prevalence of (h) functional and present ARGs predicted to originate from prophage and (i) functional and present ARGs predicted to originate from plasmids. For (a), (b), (c), (d), (h), and (i), the box plots display the interquartile range (IQR) with the median indicated as a line and the mean indicated as a triangle and whiskers extending to 1.5 times the IQR. KW refers to the Kruskal-Wallis test. Significance levels are denoted by "*", "**", "***", "****", and "ns" for adjusted $P < 0.05$, $< 0.01$, $< 0.001$, $< 0.0001$, and $\geq 0.05$ (not significant) for two-sided Mann-Whitney U tests for pairwise comparisons. Gray triangles are utilized to denote mean values.

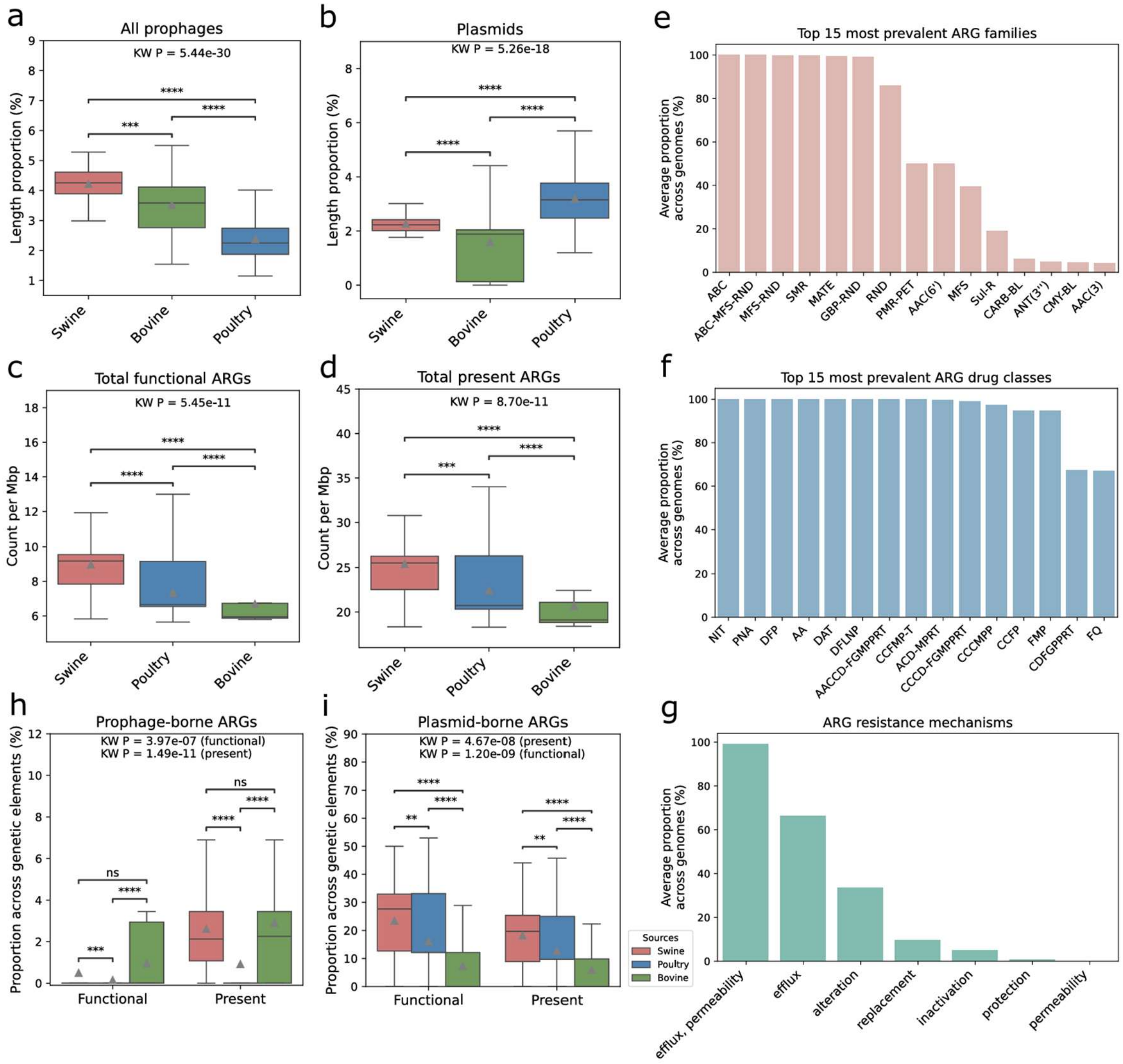


**Figure S9: AMR and MGE results for subsampled (collection year = 2021) *S.* Typhimurium genomes**. Length proportion of all identified (a) prophages and (b) plasmids identified on swine (red), wild bird (yellow), bovine (green), and poultry (blue) isolates. Total (c) putatively functional and (d) present ARGs per genome. Average proportion of the top 15 most prevalent (e) ARG families, (f) ARG drug classes, and (g) ARG resistant mechanisms. The prevalence of (h) functional and present ARGs predicted to originate from prophage and (i) functional and present ARGs predicted to originate from plasmids. For (a), (b), (c), (d), (h), and (i), the box plots display the interquartile range (IQR) with the median indicated as a line and the mean indicated as a triangle and whiskers extending to 1.5 times the IQR. KW refers to the Kruskal-Wallis test. Significance levels are denoted by "*", "**", "***", "****", and "ns" for adjusted P < 0.05, < 0.01, < 0.001, < 0.0001, and ≥0.05 (not significant) for two-sided Mann-Whitney U tests for pairwise comparisons. Gray triangles are utilized to denote mean values.

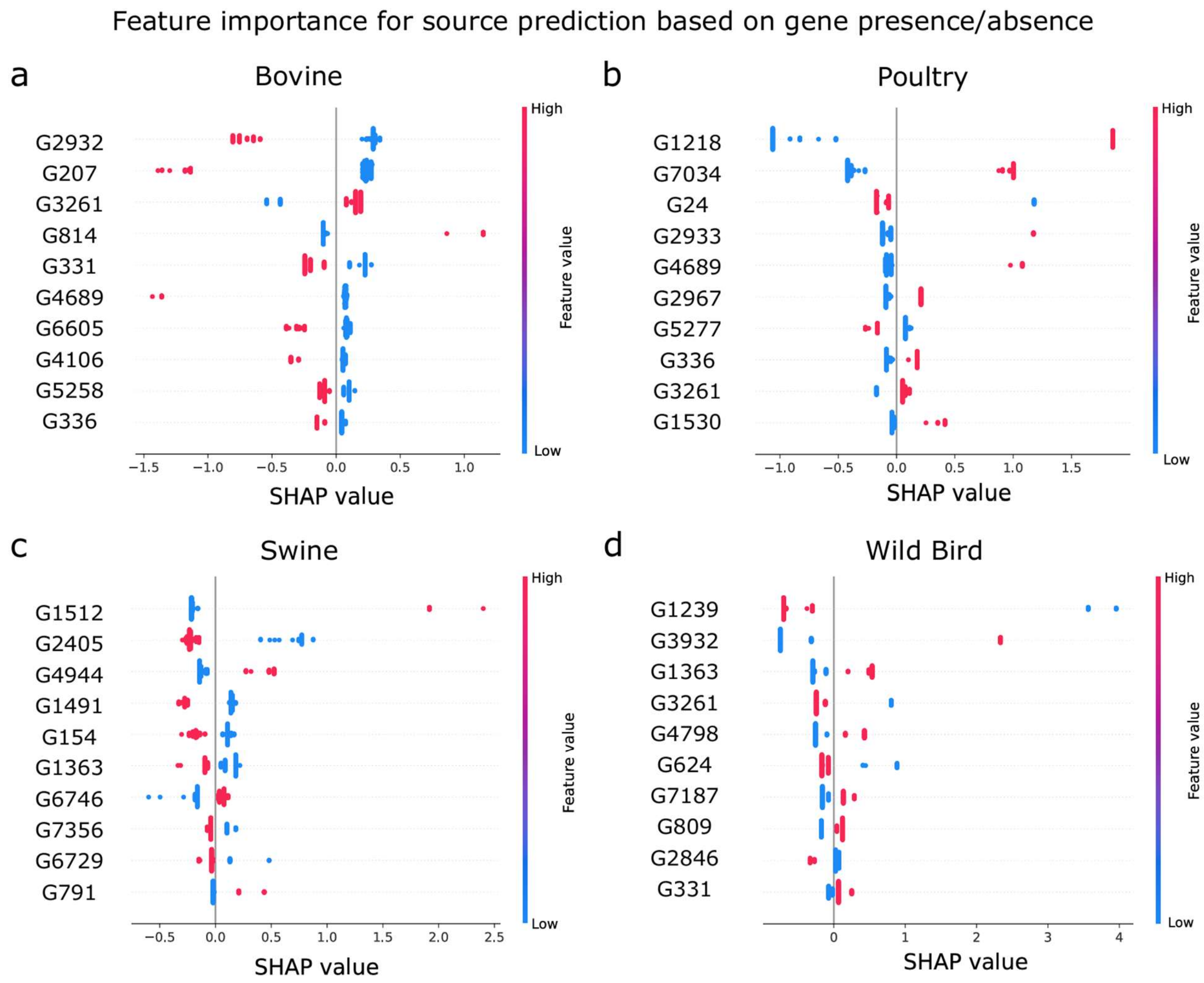


**Figure S10: Predictive genes for source attribution in *S*. Typhimurium.** The top ten most predictive genes for (a) bovine, (b) poultry, (c) swine, and (d) wild bird origins (SHAP-based; X axis), sorted by descending importance. The functional annotations of the predictive genes are provided in **Supplementary Table 5**.

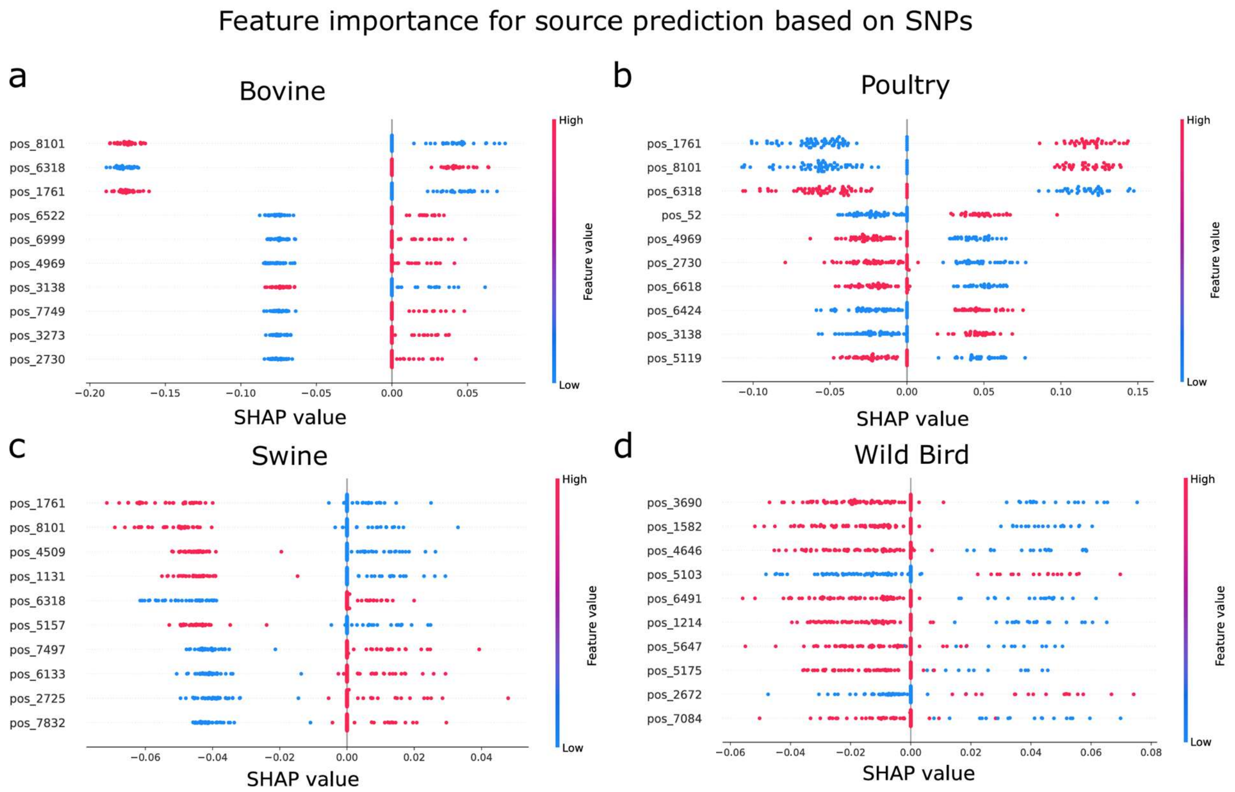


**Figure S11: Predictive SNPs for source attribution in *S*. Typhimurium.** The top ten most predictive SNPs for (a) bovine, (b) poultry, (c) swine, and (d) wild bird origins (SHAP-based; X axis), sorted by descending importance. The functional annotations of the predictive SNPs are provided in **Supplementary Table 6**.

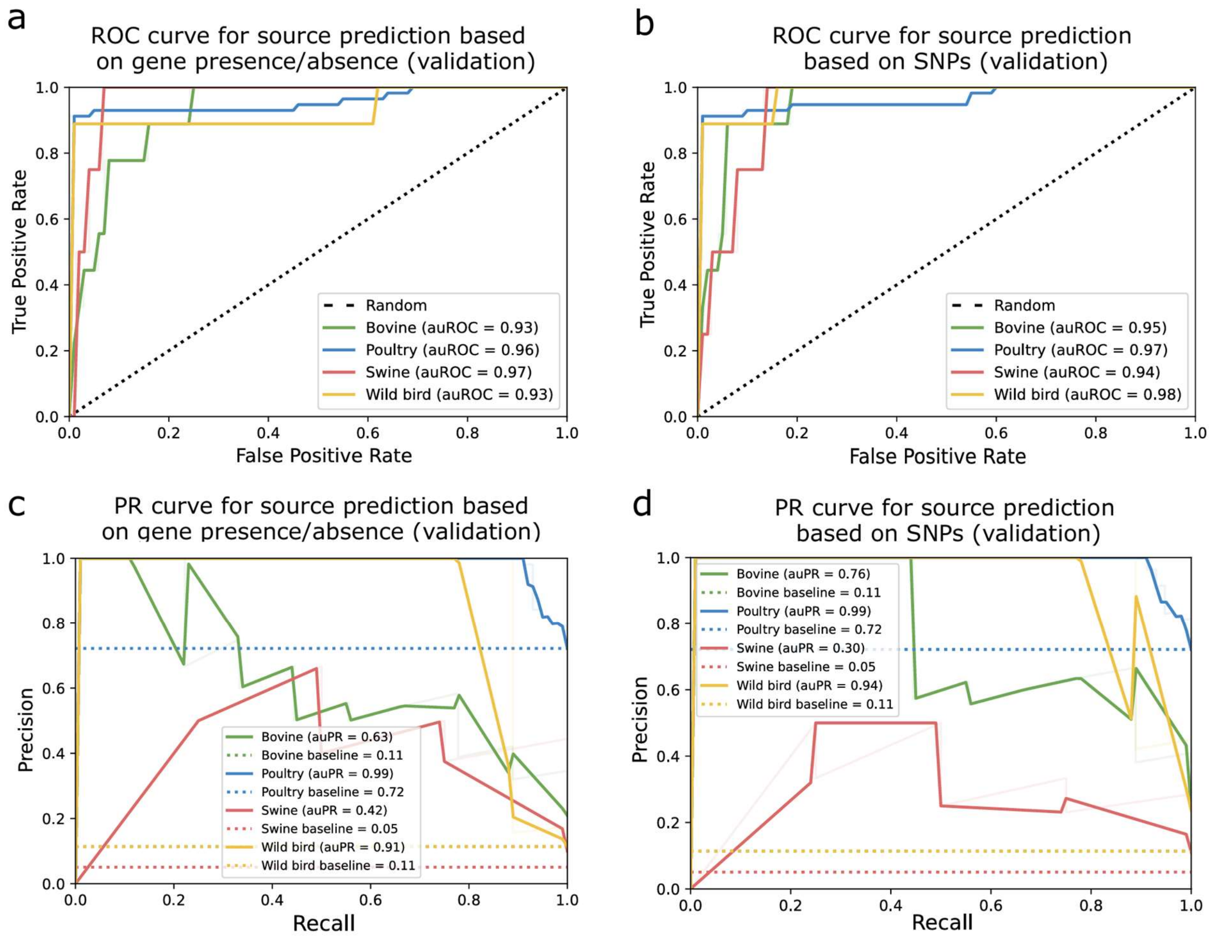


**Figure S12: Validation of machine learning models trained on gene presence/absence and core SNP profiles of *S.* Typhimurium for isolation source prediction.** Receiver operating characteristic (ROC) curves for (a) the prediction of isolation sources from gene presence/absence profiles using a lightGBM model and (b) for the prediction of isolation sources from core SNPs using a SVM model. The dark color line on the ROC curve denotes random chance accuracy. Precision-recall (PR) curves for (c) the prediction of isolation sources from gene presence/absence profiles using a lightGBM model and (d) for the prediction of isolation sources from core SNPs using a SVM model. The colored dotted lines on the PR curves are class prevalence baselines. For (a), (b), (c), and (d), each curve reflects one evaluation using test data, repeated 10 times, denoted by light color lines. The dotted lines on the PR curves are class prevalence baselines.
